# Independent genetic mapping experiments identify diverse molecular determinants of host adaptation in a generalist herbivore

**DOI:** 10.1101/2024.07.05.602111

**Authors:** Ernesto Villacis-Perez, Femke De Graeve, Berdien De Beer, Seham Ali Alshami, Rick de Jong, Tim De Meyer, Thomas Van Leeuwen

## Abstract

Interactions between plants and herbivores promote evolutionary change. Studying the evolution of herbivore mechanisms aimed to cope with different host plant species is a critical intersection between evolutionary biology and sustainable pest management. Generalist herbivores are of particular interest, as hybridization between genetically distinct populations can increase the standing genetic variation and therefore the adaptive potential of the species. *Tetranychus urticae* is a generalist arthropod known for its adaptive potential, evidenced in its immense host range and ability to develop metabolic resistance to xenobiotics. However, the molecular underpinnings associated with the potential of host adaptation and the consequences of host adaptation in this and many other pests remain elusive. Here, we use two independent, empirical approaches to identify and map the genetic basis of host plant performance and adaptation in genetically distinct populations of *T. urticae*. In the first approach, we subject a genetically diverse mite population to tomato selection and map genomic regions linked to the phenotypic evolution of increased reproductive performance. In the second approach, we map genomic regions responsible for performance on tomato by comparing the genomes of pooled individuals from a F2 backcross between populations with high and low reproductive performance. Both approaches revealed specific and shared genomic regions associated with host plant performance and adaptation and key candidate genes were identified. Our findings highlight the power of spider mite genetic approaches to identify the complex genetic basis of host adaptation in a generalist herbivore.

## 1. Introduction

Antagonistic interactions between plants and herbivores are motors of evolutionary change. Herbivory pressures plants to evolve defence mechanisms, which in turn pressure herbivores to evolve counteradaptations (Agrawal & Zhang, 2021; Futuyma & Agrawal, 2009). In agricultural ecosystems, herbivores face additional pressure from pesticides and resistant plant varieties, and readily evolve adaptations to overcome these challenges (Bunga et al., 2024; De Rouck et al., 2023). Understanding the factors underlying the adaptive potential of herbivores and the consequences of adaptation across different levels of biological organization is crucial to building sustainable crop management strategies (European Commission, 2021; Van Leeuwen et al., 2020).

Plants produce an array of chemical and structural defences aimed to repel herbivores, hinder their development, or attract their natural enemies (Aljbory & Chen, 2018; Erb & Reymond, 2019; Snoeck et al., 2022). Closely related plant taxa often share similar defensive strategies (but see e.g. Petschenka et al., 2017), and thus herbivore counteradaptations are largely dependent on their host range (Ali & Agrawal, 2012). Specialists can cope with few hosts, but do so very efficiently by evolving insensitivity to toxins, by detoxifying them, or by suppressing their production (Aggarwal et al., 2014; Blaazer et al., 2018; Herfurth et al., 2017; Shih et al., 2023; Wang et al., 2023; Zhao et al., 2015). In contrast, generalists exploit a wide range of hosts, requiring flexibility in their mechanisms of host use (Grbić et al., 2011). Plasticity in the molecular mechanisms that regulate genes involved in host use can enable generalists to cope with multiple classes of phytochemicals (Ji et al., 2023; Pym et al., 2019; Snoeck et al., 2018; Vandenhole et al., 2021). While generalists and specialists largely employ similar, but not identical mechanisms to exploit their hosts (Ali & Agrawal, 2012; Bruinsma et al., 2023; Bui et al., 2018), it is not well understood how selection shapes the diversity of mechanisms involved in host use in generalists, or whether genetic changes due to selection impact the host range of locally adapted populations.

Occurring on over 1100 plant species across over 130 plant families (Migeon et al., 2024), *Tetranychus urticae*, the two-spotted spider mite, is growing as a model to study adaptive evolution. While often characterised as an extreme generalist, genetic and phenotypic variation related to host use occurs between and within its populations (L. Chen et al., 2023; Kant et al., 2008; Mortier et al., 2021; Villacis-Perez et al., 2021; Xue et al., 2023). *T. urticae* tops the list of arthropods with the most cases of resistance to the largest number of pesticide classes (De Rouck et al., 2023). Its genome shows a remarkable expansion of genes potentially involved in host use, including detoxification families such as cytochrome P450s (P450) and glutathione-S-transferases (GSTs), transporter families such as ATP-synthase binding cassette (ABC) genes, major facilitator superfamily (MFS) transporters (Dermauw et al., 2013; Grbić et al., 2011; Schlachter et al., 2019), as well as an extreme expansion of chemosensory receptor families (Ngoc et al., 2016). In addition, gene families acquired ancestrally by horizontal gene transfer aid in xenobiotic detoxification, such as intradiol ring cleavage dioxygenases (DOGs), UDP-glycosyltransferases (UGTs) and a β-cyanoalanine synthase (CAS) (Njiru et al., 2022; Snoeck et al., 2019; Wybouw et al., 2014, 2018)

Spider mites are excellent laboratory organisms. With a short generation time of 12-14 days under laboratory conditions, populations experimentally isolated on challenging hosts show clear adaptation patterns within 5 to 25 generations (Belliure et al., 2010; Cazaux et al., 2014). Genetically distinct populations often show no signs of reproductive isolation, allowing to create hybrid lines in which parental alleles can be traced and recombination is even (Ji et al., 2023; Xue et al., 2023). In addition, a well-annotated, chromosome-level assembled genome facilitates the integration of bioassays with genomic tools to identify causal genes within quantitative trait loci (QTLs) (Grbić et al., 2011; Kurlovs et al., 2019; Wybouw et al., 2019). For example, bulked segregant analyses (BSA) paired with whole genome sequencing have successfully mapped QTL regions underlying resistance to multiple classes of acaricides (De Beer et al., 2022; Vandenhole et al., 2023; Villacis-Perez et al., 2023; Wybouw et al., 2019). However, the identification of loci that respond to selection imposed by host plants remains elusive.

In this study, we conduct two complementary experiments to identify the genetic determinants of adaptation and performance on a host previously shown to be challenging for this species: tomato (*Solanum lycopersicum*; Kant et al., 2008). In the first approach (referred to as ‘Experimental Evolution’), genetic determinants are identified through an adaptation experiment, where we create a quasi-panmictic spider mite population from 6 genetically distinct parents and subject it to selection by an experimental shift from bean to tomato. We quantify evidence of adaptation across replicated populations after 20 generations of selection, map QTLs and evaluate the impact of adaptation on gene expression. In a second approach (referred to as ‘F2 Backcross’), genetic determinants are identified in an F2 population using two genetically unrelated populations with clear phenotypic differences in performance on tomato. We identify genomic regions responsible for tomato performance by comparing genomes from pooled F2 individuals with high and low reproductive performance on tomato. Our analyses revealed specific and shared genomic regions between both approaches. We discuss the implications of these findings for the evolution of host use mechanisms in generalist herbivores.

## 2. Methods

### 2.1 Plants

Common bean (*Phaseolus vulgaris* cv. Prelude or Speedy), tomato (*Solanum lycopersicum* cv. Moneymaker, cv. Castlemart and cv. def-1), maize (*Zea mays* cv. Ronaldinho) and sweet pepper (*Capsicum annum* cv. Spider F1) plants were grown under standard conditions (25 ± 1°C, 60% relative humidity [RH], 16h:8h light:dark [L:D] photoperiod,) in the greenhouse of the University of Amsterdam. Plants 2-3 weeks old (bean and pepper) and 4-5 weeks old (tomato and maize) were used for experiments. Wild honeysuckle (*Lonicera periclymenum*) leaves were obtained from potted plants that were originally acquired from a local garden shop.

### 2.2 Experimental Evolution approach

To map loci associated with host adaptation, we generated a genetically diverse, quasi-panmictic population by crossing six iso-female lines of *T. urticae* with green body colouration established previously (Table 1; Xue et al., 2020). Lines were maintained in the laboratory under standard conditions on detached bean leaves placed on wet cotton wool to prevent cross-contamination. To address the potential issue of partial reproductive incompatibility previously reported for *T. urticae* (de Boer & Veerman, 1983; Overmeer & Van Zon, 1976), the six lines were divided into three blocks, each of which was comprised by ten virgin females from one line, ten virgin females from a second line, ten males from a third line and ten males from a fourth line. Then, 20 virgin females from the resulting F1 generation from each block were crossed with 20 males of the remaining two lines (i.e., fifth and sixth line). The F2 offspring from these crosses were transferred together to multiple bean plants and the population expanded for approximately 4 to 5 generations before further experiments. This population is henceforth referred to as the ‘Mix’.

**Table 1.** Parental *Tetranychus urticae* lines used to create the experimental ‘Mix’ population. Name and laboratory ID* of the six iso-female expansion lines derived from field populations from different host plants and geographical locations.

| Name | Lab ID* | HOST | LOCATION |
| --- | --- | --- | --- |
| BE | BE-B8.17 | Potato | Belgium |
| DE | DE-B8.13 | Potato | Germany |
| ES | ES-B8.19 | Strawberry | Spain |
| NL | NL-B8.9 | Strawberry | Netherlands |
| PL | POL-G | Tomato | Poland |
| UK | UK-B7.6 | Cucumber | United Kingdom |

\* Laboratory ID follows the nomenclature of the lines as in Xue et al., 2020

We subjected the Mix to an experimental evolution assay using tomato as a challenging host, which the Mix had not experienced before. First, ten samples of ∼350 mated females were transferred from the expanded Mix population to mite-proof cages with bean plants into a greenhouse set to standard conditions. After two generations (∼28 days), a highly infested leaf with >500 individuals was transferred from each bean cage to a paired sibling cage with a tomato plant. Thus, each of the ten replicates created consisted of a bean population paired to a sister population on tomato. Populations on both plants were allowed to develop in large numbers to prevent bottlenecks, and new plants were added to the cages before the plants were overtaken, approximately every 12 days. To prevent cross-contamination, cages were isolated from each other by placing them on top of a table with soapy water. Twenty generations after infesting the tomato cages, approximately 800 adult females from each of the bean and tomato populations were collected, flash frozen in liquid nitrogen and stored at -80°C before DNA extraction, whole genome sequencing, subsequent quality control and variant calling (Supplementary Note 1).

We also assessed the changes in gene expression upon experimental evolution. To do so, we transferred ∼150 females from five paired replicates on bean and tomato to detached bean leaves and allowed them to develop for one generation. Thus, mites from tomato were reverted to bean for one generation and synchronised in age with the bean populations. 150 adult females (2 ± 1 days old) from each of these synchronised cohorts were collected and transferred to either a bean or a tomato plant in the laboratory. After 24 hours of feeding, 100 alive individuals per plant were collected for each sample. A total of 20 samples were obtained, comprising five samples from the bean mites feeding on bean, five samples from the bean mites feeding on tomato, as well as their paired tomato sisters on both hosts. Samples were flash frozen in liquid nitrogen and stored at -80°C until RNA extraction, sequencing, differential gene expression analysis and gene ontogeny (GO) enrichment analysis (Supplementary Note 1).

#### 2.2.1 Phenotyping of fitness traits

To infer the extent of variation in traits associated with host use in the parental lines, we measured the reproductive output of individual females, juvenile survival, and juvenile development until adulthood, on bean and tomato. The same fitness traits were measured for the Mix after its creation and subsequent expansion. We also quantified these fitness traits on the populations subjected to tomato selection 20 generations after the shift from bean. To discern between genetic evolution and acclimatisation or potential maternal effects, we defined four treatments: 1) mites from bean; 2) mites from tomato; 3) tomato mites reverted for one generation to bean; 4) tomato mites reverted for two generations to bean. Experimental set-ups to measure each trait and data analyses are specified in Supplementary Note 2.

#### 2.2.2 Genomic scans

To map loci associated with tomato adaptation, we assessed differences in allele counts between bean and tomato mites by fitting a binomial generalized linear model (GLM) to each locus that passed the filtering criteria (Supplementary Note 1) using the *glm* function in R (v4.3.1), with selection host and experimental replicate as fixed effects. Significance of the effect of selection host on the allele counts was determined through a likelihood ratio test (LRT) using the *anova* function. To address potential issues of underestimated p-values due to overdispersion, we followed the methodology described by Hoedjes et al., 2019 for FDR correction (Supplementary Note 3). SNPs were considered significant if the mean FDR was below 0.01. Similar to the strategy described by Zhang & Panthee, 2020, we subsequently used a sliding window approach (window size of 150 kb, step size of 15 kb) to identify regions likely under selection. For each window, we calculated the fraction of significant SNPs (FDR < 0.01) to the total number of SNPs within the window. The idea behind this approach is that regions harbouring genes under selection, and those in close proximity due to linkage, would exhibit a higher fraction of significant SNPs.

A genome-wide threshold for significance of windows was calculated based on random sampling of an equivalent number of SNPs as the genome-wide average count per window. For this random window, we computed the fraction of significant SNPs, and repeated this process 10,000 times. The 99^th^ percentile of all fractions served as the threshold for identifying significant windows. To enhance visualization, we smoothed the observed averages using a Savitzk-Golay filter from the Python (v3.10.6) package *scipy* (v1.11.4) (Virtanen et al., 2020). We used a modified version of the BSA-scan method available at https://github.com/rmclarklab/BSA (Wybouw et al., 2019) to verify our findings (Supplementary Note 3).

### 2.3 F2 backcross approach

To map loci associated with reproductive performance on tomato, we used two laboratory *T. urticae* lines with red body colouration: VR-BE, a field line obtained from tomato plants (Njiru et al., 2023), and Pol-R, an iso-female line developed previously in the laboratory. Lines were kept under standard conditions on detached bean leaves surrounded by wet-cotton wool. Sixty virgin females from Pol-R were crossed with 30 males from VR-BE, producing an F1 hybrid population on detached bean leaves. Approximately 280 virgin F1 females were then collected and split into two groups. The first group was used to phenotype the reproductive performance of the F1 hybrids and the parental lines. The second group was used to create an F2 backcross. F1 virgin females were isolated on a new bean leaf with approximately 100 males from the Pol-R line and allowed to lay eggs for two days, after which all mites alive were transferred to a new detached leaf, and so on every two days. Thus, we created large age-synchronised cohorts of F2 individuals for further phenotyping assays. Several Eppendorf tubes containing pools of 150 – 200 random, non-phenotyped F2 females from these cohorts were flash frozen and stored at -80°C, and served as controls for the mapping assay.

#### 2.3.1 Phenotyping of fitness traits

To characterise the genetic basis of reproductive performance, we measured the number of eggs laid per female per day using the parental lines, F1 hybrids and the F2 backcross. Age-synchronised adult females (2 ± 1 day old) from the parental lines and their F1 offspring were isolated each on a 15mm (diameter) bean or tomato leaf disc surrounded by wet cotton wool (n = 32 per line, per plant). Female mortality was scored daily, and the number of eggs laid per female was scored after four days. Oviposition was corrected for mortality. To assess the reproductive performance of the F2 backcross, age-synchronised females (2 ± 1 day old) from the F2 cohorts were transferred to tomato leaf discs, and their mortality-corrected oviposition was assessed as described above. Five consecutive cohorts were scored for oviposition on tomato, for a total of n = 933 females. After scoring survival on the fourth day, each individual female was collected in a PCR strip tube and stored at -80°C. After female performance was calculated, we selected individuals based on high and low performance groups. Approximately 150 females were pooled into a single Eppendorf tube per group over a bed of dry ice to prevent degradation and re-stored at -80°C until DNA extraction, whole-genome sequencing, subsequent quality control and variant calling, along with the non-phenotyped F2 pooled samples (Supplementary Note 1).

#### 2.3.2 Genomic scans

We employed the same method described in section 2.2.2 to detect loci associated with reproductive performance in the F2 approach, with some adjustments. The window and step size were set to 250 kb and 25 kb, respectively. Because we pooled individuals into single samples, we scanned the genome solely for AF differences between the high-performing and the control samples, as well as between the low-performing and the control samples. The control sample consisted of 150 random F2 females that were not phenotyped. We used a Fisher exact test to identify differences in allele counts between the high-performing and the control samples, as well as between the low-performing and the control samples, on a per-locus basis, following the standard approach described by Zhang & Panthee, 2020. We calculated average fractions of significant SNPs (p-value < 0.01) per window, and the genome-wide threshold for window significance using the same procedure as in section 2.2.2.

## 3. Results

### 3.1 Experimental Evolution approach

To map loci associated with host adaptation using an experimental evolution approach, we crossed six parental lines (Table 1) in order to maximise the chances that genetic and phenotypic variation relevant to host adaptation would be captured in their offspring. We exposed this offspring, the ‘Mix’ population, to a host shift from bean to tomato and measured genomic, transcriptomic, and phenotypic changes after 20 generations.

#### 3.1.1 Genetic and phenotypic variation across parental lines and the Mix

To understand the extent of genetic variation across lines, we sequenced the genomes of the parental lines and the Mix upon its creation. The parental lines yielded between 27 to 41 million raw 150 bp paired-end Illumina reads, while the Mix generated 110 million 75 bp paired-end reads. To assess genetic differentiation between lines, we conducted a PCA using 442,394 high-quality SNPs distributed across the genome (Supplementary Note 1). The first four principal components (PC) explained 26.7%, 21.8%, 17.7%, and 17.2% of the variance, respectively (Figure 1). The six parental lines were distinct from each other, with no apparent clustering suggestive of close relatedness. Although lines DE, BE, and NL clustered closely together on PC1 and PC2, they were distinctly separated on PC3 and PC4.

**Figure 1.**
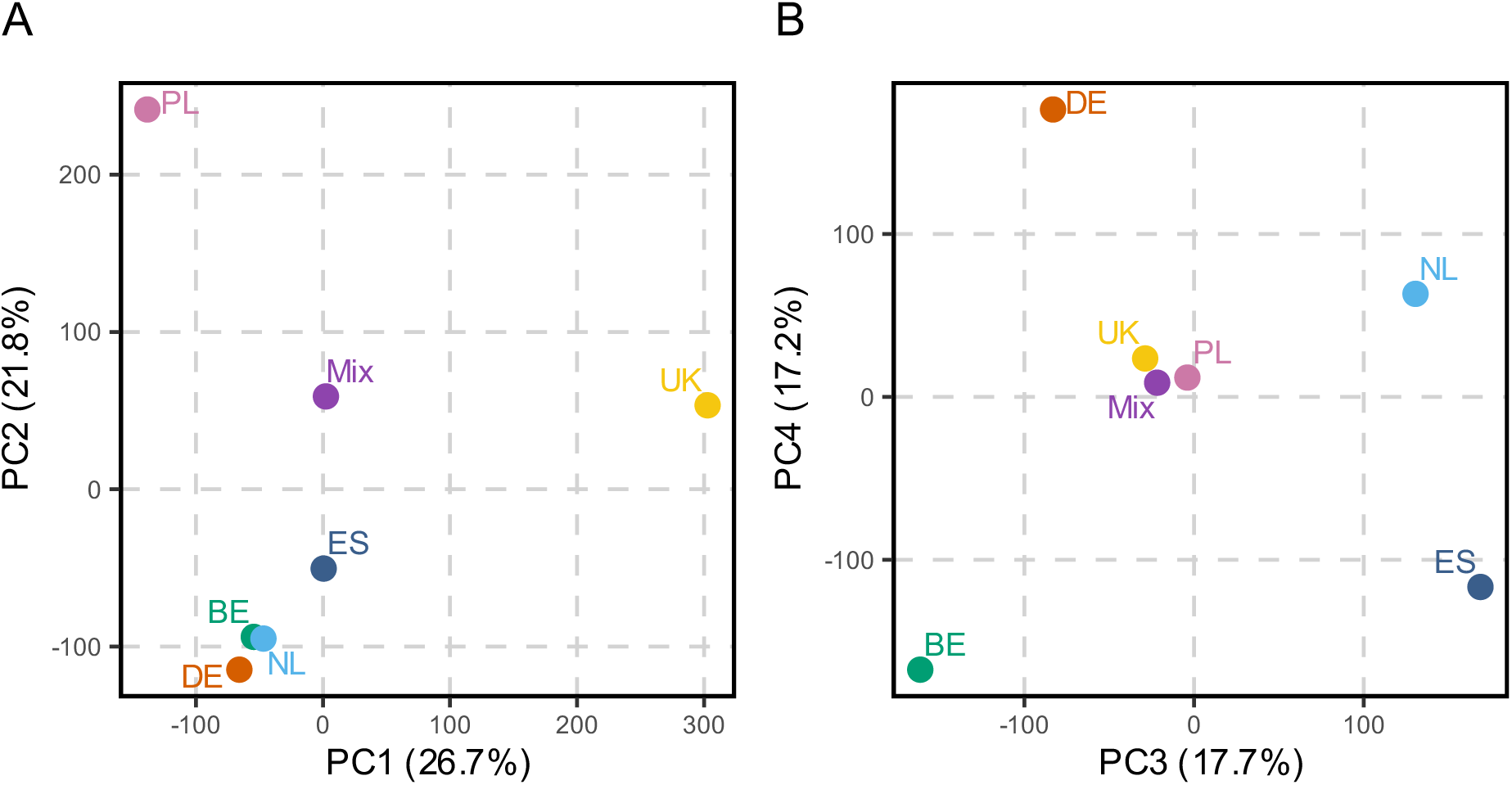
Genetic variation among the parental lines and the Mix. PCA of the parental populations and the Mix using genome-wide allele frequency data at polymorphic sites. Each point represents a strain, labeled with its corresponding name. The percentage of explained variance is shown in parentheses for all principal components. Panels show PC1 vs PC2 (A) and PC3 vs PC4 (B).

Tracking homozygous alleles unique to each line revealed uneven contributions of the parental lines to the genomic profile of the Mix. Lines PL and ES had the largest differences, contributing approximately with 40% and 2% to the profile, respectively. The UK line contributed with ∼22%, NL with ∼9%, BE with ∼14% and DE with ∼13%. This imbalance is evident along PC3 and PC4, where the Mix clusters most closely with PL and UK, the two most represented parental lines in the Mix (Figure 1B).

We found significant variation in reproductive output, juvenile survival, and developmental time among the parental lines, on bean and tomato (Supplementary Figures 1-2). Most lines laid significantly more eggs on bean compared to tomato (average across lines: 5.73 ± 0.20 (mean across lines ± standard error of the mean) on bean vs. 4.08 ± 0.18 on tomato, except for line NL, which had a relatively low performance on both hosts (Supplementary Figure 1A-F; Supplementary Table 1). Comparisons within plant species showed significant variation in egg numbers across lines for bean and tomato (Supplementary Figure 2A-B; Supplementary Table 2). Some of the lines (e.g., lines BE, UK, DE) harboured individuals that laid twice as many eggs as the overall average (>10 eggs on bean and > 8 eggs on tomato), but individuals with a low performance (<2 eggs per day) were overall common across lines (Figure 1A-F). Juvenile survival differed significantly across lines in both bean and tomato (Supplementary Figure 2C-D; Supplementary Table 3A). For most lines, the probability of juvenile survival was significantly lower on tomato than on bean, but this was highly dependent on the line (Figure 1G-L; Supplementary Figure 2C-D; Supplementary Table 3A). For example, the survival of juveniles from lines BE, PL and UK did not differ between bean and tomato (Figure 1G, K, L; Supplementary Table 3B). However, while for lines BE and UK the survival probability progressively decreased until reaching ∼50-65% by the last time point on day 13, the survival of line PL barely decreased over time and was >95% throughout the experiment. Juvenile survival was significantly different for lines DE, ES and NL between plant species, with survival on tomato being significantly lower than on bean (Figure 1H, I, J; Supplementary Table 3B). Significant variation in developmental time was found across lines. Most lines had a significantly slower development on tomato compared to bean (Supplementary Table 3D), with juveniles from lines DE and ES never developing to adulthood during the experimental period, and lines BE, NL and UK reaching a probability between 25-50% of becoming adults. In strike contrast, line PL developed as quickly on bean as it did on tomato, with over 25% of individuals reaching adulthood by day 9, over 70% by day 11 and >99% reaching adulthood on both plants by the last time point (Figure 1Q; Supplementary Figure 1M-R; Supplementary Figure 2E-F; Supplementary Table 3C). In summary, the parental lines feature clear genetic and phenotypic differences from each other.

The Mix line had a higher female reproductive output on bean than on tomato (Supplementary Figure 3A; Supplementary Table 2), with more eggs laid on both plants compared to any parental line (bean: 8.52 ± 0.21, tomato: 5.44 ± 0.32; Supplementary Figure 1A-F). Juvenile survival was high on both plants (80-90%), but unlike the parental lines, it was significantly lower on bean than on tomato (Supplementary Figure 3B). The developmental time to adulthood was slower on tomato than on bean, but all surviving juveniles reached adulthood by the last time point on both plants (Supplementary Figure 3C).

#### 3.1.2 Tomato selection yields high reproductive performance across hosts

To discern between genetic evolution and acclimatisation or potential maternal effects, we defined four treatments: 1) mites from bean; 2) mites from tomato; 3) tomato mites reverted for one generation to bean; 4) tomato mites reverted for two generations to bean.

##### 3.1.2.1 Phenotypic evidence of tomato adaptation

After 20 generations, the bean replicates maintained a similar reproductive output on bean and tomato as when the Mix was created, with significantly more eggs laid on bean (7.94 ± 0.17) than on tomato (5.71 ± 0.16; Figure 2A; Supplementary Figure 3; Supplementary Table 4). The mites from tomato had the largest reproductive output across all treatments on tomato (9.83 ± 0.11). Reverting the tomato mites for either one (7.42 ± 0.13) or two generations (7.08 ± 0.13) to bean decreased their reproductive output on tomato, but their output was still significantly higher than the control bean mites, providing evidence of genetic adaptation to tomato (Figure 2; Supplementary Table 4). Mites from tomato also laid significantly more eggs on bean than the bean mites (Figure 2B; Supplementary Table 4). The reproductive output of tomato mites after being reverted to bean for one generation was the largest across treatments on bean (9.45 ± 0.13). After being reverted for two generations to bean, the number of eggs laid per female decreased again, reaching a midpoint between the bean mites and the tomato mites (8.35 ± 0.15; Figure 2C-D; Supplementary Table 4).

**Figure 2.**
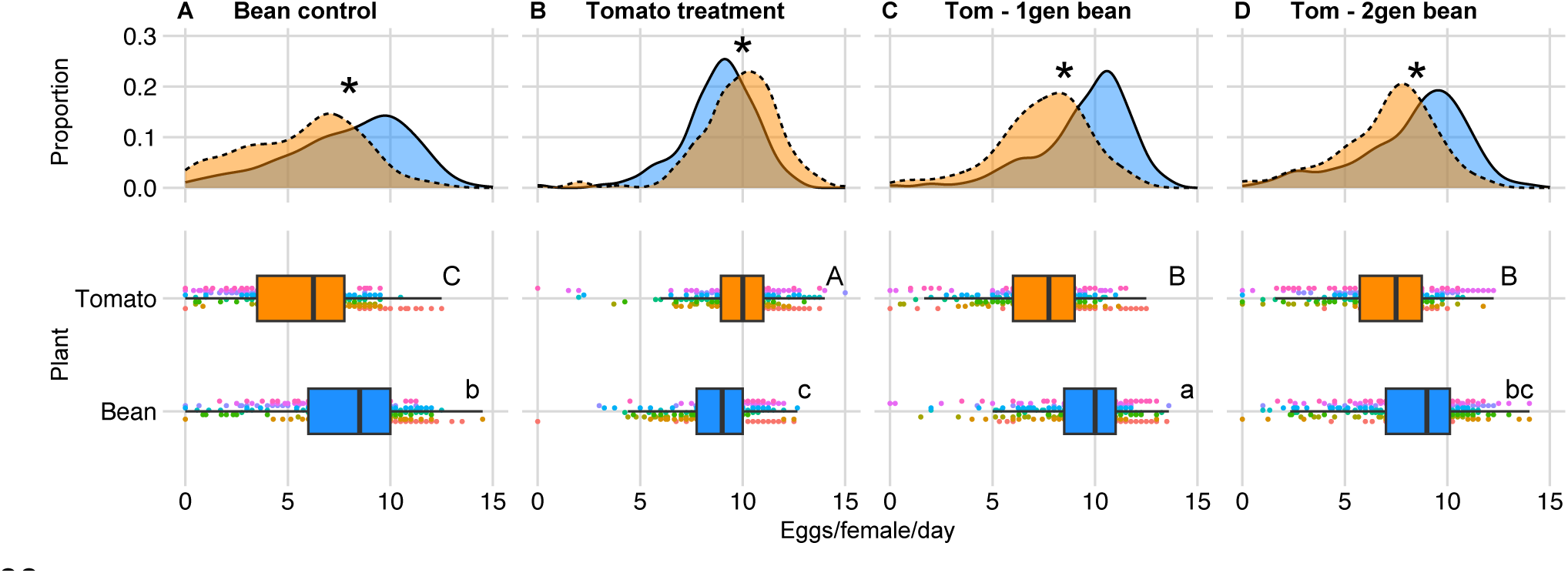
Reproductive output of adult females after 20 generations of experimental evolution. The number of eggs laid per female per day alive was measured on bean and tomato for the un-selected control bean lines (A), the tomato-evolved lines (B), and the tomato-evolved lines after being reverted to bean for either one (C) or two (D) generations. The distributions on the top panels show the proportion of females (y-axes) that laid a certain number of eggs (x-axes) on tomato (in orange and dotted lines) and bean (in blue and solid lines), while the boxplots in the bottom panels show the individual data points coloured by replicate (Replicates 1-10; circles), the median (black line within the interquartile box), and the bottom and top 25% quartiles of the data values (whiskers) on tomato (in orange) and bean (in blue). Stars on the top panels show a significant difference in the number of eggs laid between bean and tomato according to a linear mixed model fitted to the data per panel. Uppercase letters in the lower panels show significant differences on the number of eggs laid per female on tomato across panels A-D, and lowercase letters show differences across panels for bean, according to linear mixed models applied to either the tomato or the bean data. Per boxplot, n = 313-320 females.

Juvenile survival on both bean and tomato was similar to the original Mix (∼75-85% by day 13), except for a decrease of the survival of the tomato mites reverted to bean for two generations on bean (Supplementary Figure 4A-B). The probability of reaching adulthood varied across treatments on both plants (Supplementary Figure 4C-D). On bean, tomato mites and two-generation reverted mites reached adulthood faster than controls and than the mites reverted to bean for one generation, but all individuals reached adulthood by the last time point. On tomato, individuals from tomato and bean-reverted treatments reached adulthood quicker than the bean mites (Supplementary Figure 4D).

To assess whether tomato selection influenced fitness across other host species, we scored the reproductive output of age-synchronised females from a subset of three bean replicates, their paired tomato sisters, and tomato mites reverted to bean for two generations, across multiple host plants (Supplementary Note 2). Mites from tomato and groups reverted to bean laid more eggs than bean mites on bean, tomato, and maize, but not on honeysuckle or sweet pepper (Supplementary Figure 5A). Additionally, we assessed reproductive performance on tomato varieties differing in their jasmonic acid defence profile and found that while performance did not differ across varieties, it highly differed between treatments, with the mites from tomato laying the largest number eggs compared to bean mites and tomato mites reverted to bean (Supplementary Figure 5B).

##### 3.1.2.2 Genomic and transcriptomic evidence of tomato adaptation

After 20 generations of selection, we sequenced the genomes of five paired replicates per host. Between 33 and 36 million Illumina reads were generated and mapped to the ‘London’ reference genome for subsequent variant calling. The PCA plot constructed based on 1,109,452 genomic SNPs showed clear evidence of genomic changes associated with tomato selection. The tomato and the bean replicates clustered separately in two groups along PC1, which explained 31.1% of the variation in the SNP dataset (Figure 3A). The tomato replicates were more dispersed as compared to the bean replicates along PC2, which explained 17.2% of the variation. We calculated the genome-wide average heterozygosity proportion between the tomato and bean replicates. On average (mean ± SEM), tomato replicates had significantly lower heterozygosity than the bean replicates (bean: 0.89 ± 0.002 vs. tomato: 0.85 ± 0.015; *t* = 6.11, df = 4, p = 0.004).

**Figure 3.**
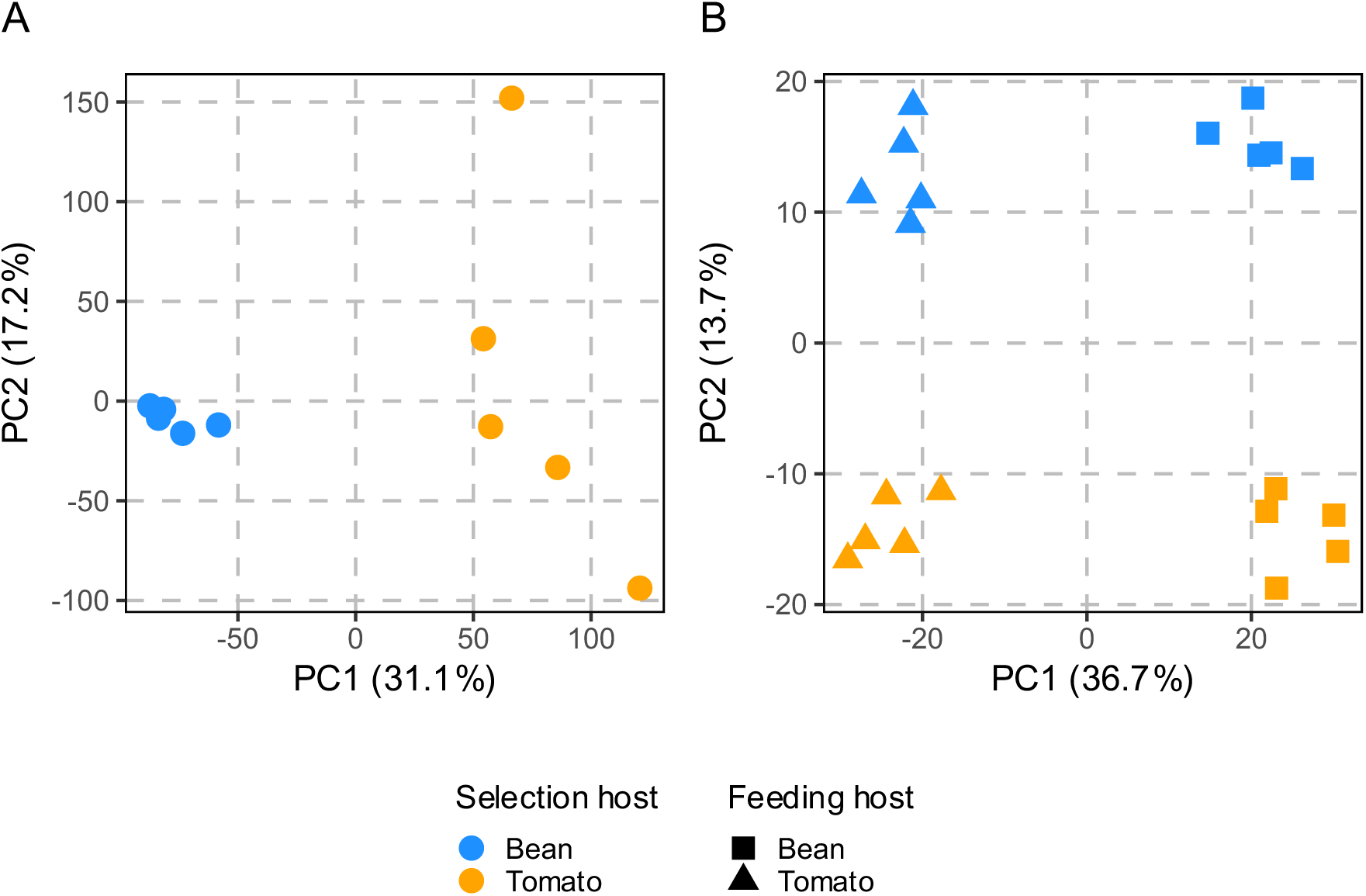
Genomic and transcriptomic profiles of lines after 20 generations of experimental evolution. A) Genomic PCA of the unselected bean control and tomato-selected populations using genome-wide allele frequency data at polymorphic sites. B) Transcriptomic PCA of the unselected bean control populations and the tomato-selected populations, feeding either on bean or tomato, using genome-wide normalized gene counts. Explained variance percentages are given in parentheses.

The PCA plot constructed on the gene expression profiles of the same five paired replicates showed a clear distinction between both selection and feeding host plant. Along the first two components, which explained 36.7% and 13.7% of the variation in the expression data, the replicates clustered in four distinct groups (Figure 3B). The clusters grouped along PC1 separated the treatments based on feeding host, while PC2 separated the replicates into either experimental evolution host.

##### 3.1.2.3 Loci associated with tomato adaptation

To identify loci responding to selection, we calculated the fraction of significant SNPs (FDR < 0.01) along sliding genomic windows. The significance threshold for windows in this method was calculated through permutation of windows with a random selection of loci across the genome (see section 2.2.2; Supplementary Note 3). The clearest peaks exceeding the genome-wide significance threshold were located on chromosome 3: QTL1 at 7-8 Mb (cumulative genomic position at ∼70 Mb) and QTL2 at 15.5 Mb (cumulative position at ∼78 Mb) (Figure 4A). These peaks were verified by the AF difference plots obtained from the alternative BSA analysis strategy (Supplementary Note 3; Supplementary Figure 7A). We analysed these regions further for candidate genes potentially linked to tomato adaptation. Apart from these two large peaks, multiple, although smaller, significant peaks were detected on chromosomes 1 and 3, but only two minor peaks on chromosome 2.

**Figure 4.**
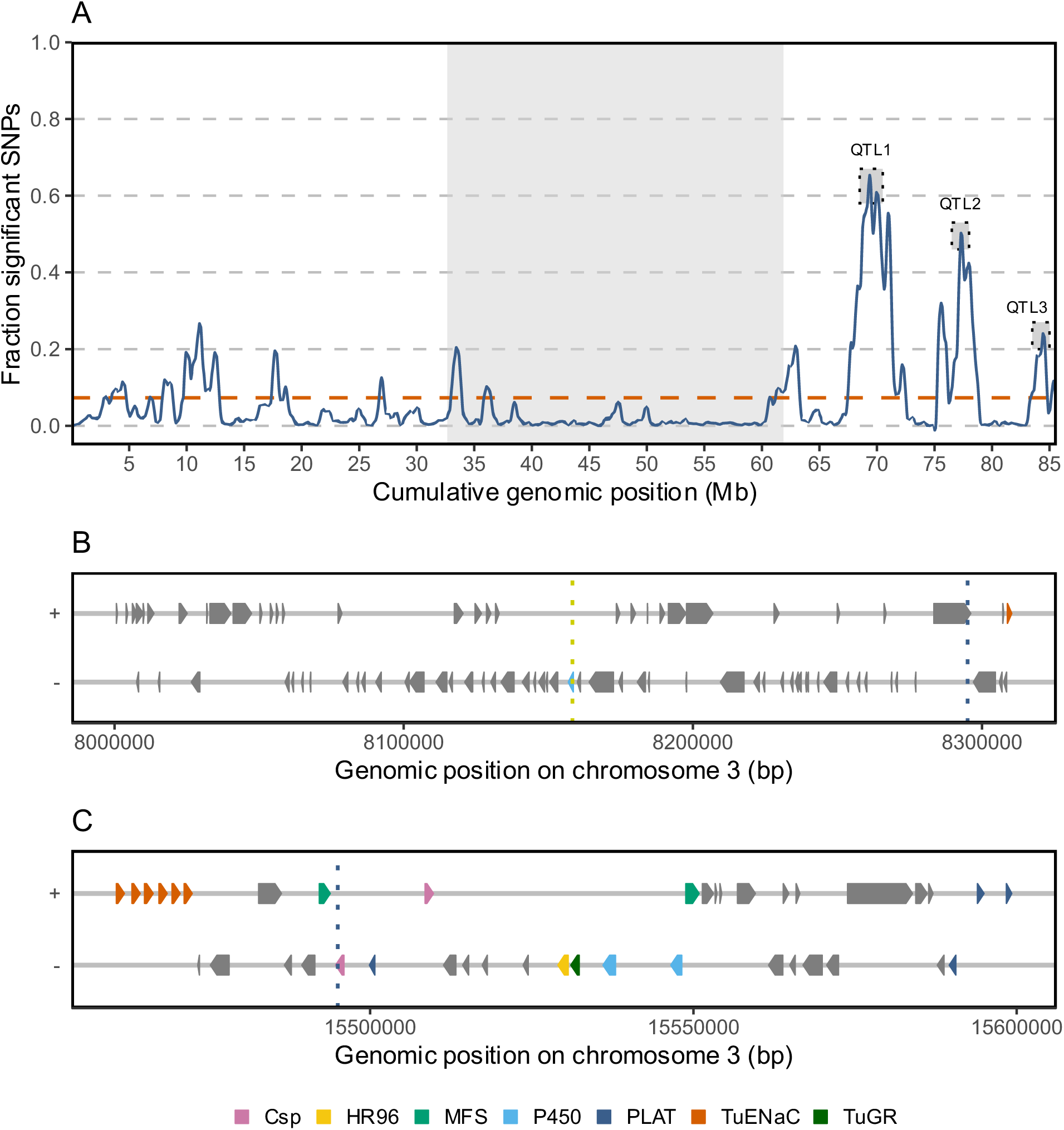
Genomic response to tomato selection after 20 generations of experimental evolution. Panel (A) showing the fraction of significant SNPs (FDR < 0.01) calculated in sliding windows. The dashed red line shows the genome-wide significance threshold for windows, determined by random permutation of windows. Chromosomes are arranged in descending order and indicated by alternating shading. Genes within selected genomic regions around QTL1 (B) and QTL2 (C) are coloured (legend) according to their class if deemed interesting, otherwise they are shown in grey (Csp: conserved secreted protein, HR96: DBD lacking nuclear hormone receptor, MFS = major facilitator superfamily, P450 = cytochrome P450, PLAT = lipase/lipoxygenase, TuENaC: degenerin/epithelial Na+ channel chemosensory receptors, TuGR = gustatory chemosensory receptor). Vertical dotted lines represent the top genomic window (blue), or the peak in frequency of PL alleles in the tomato-selected samples (yellow).

For QTL1 and QTL2, we examined whether the region could be associated with the fixation of alleles from specific parental lines by analysing the mean AF in bean and tomato replicates using unique parental marker SNPs (Supplementary Figure 6). Within QTL1 there was near fixation of PL alleles in the tomato replicates (Supplementary Figure 6A). The mean AF of unique, homozygous PL alleles in the tomato mites reached a maximum at ∼ 8.158 Mb, within a gene from the P450 family: CYP392B3. For QTL2, both homo- and heterozygous unique NL marker SNPs were examined, as the parental line was heterozygous in the considered region. We found evidence of selection towards NL alleles in this region in the tomato replicates (Supplementary Figure 6B).

For QTL1, we analysed a 300 kb region surrounding the peak of homozygous unique PL marker SNPs (Figure 4B, Supplementary Table 7). The most noteworthy candidate in this region is *tetur20g00290* (CYP392B3), with the peak of PL markers located within this gene. Additionally, we identified 13 missense variants and one high-impact variant (i.e. gained stop codon) within this gene, unique to the PL line (Supplementary Table 8). The PL alleles for all these variants were nearly fixed in the tomato replicates (AF > 0.93). We found potential evidence of an extra copy of the gene in the PL line, as suggested by DNA read coverage and heterozygosity within an otherwise homozygous parental region. Furthermore, other candidate genes in the region, including *tetur20g00360* (Short-chain dehydrogenases/reductase gene), *tetur20g00330* (Sodium/hydrogen exchanger), *tetur43g00480* (TuENaC05, degenerin/epithelial Na+ channel receptor), *tetur43g00470* (glycoside hydrolase) and *tetur43g00460* (ionotropic glutamate receptor), had unique PL missense variants, which nearly reached fixation in the tomato replicates. The transcriptomic data did not show DE for any of the candidate genes discussed above. However, five genes within the QTL1 region, *tetur20g00400* (PREDICTED: similar to Bardet-Biedl syndrome 7) and four hypothetical proteins, exhibited constitutive downregulation upon tomato adaptation.

For QTL2, which was smaller than QTL1, we analysed a 100 kb region centred around the peak window (Figure 4C, Supplementary Table 9). Alongside 11 genes from known detoxification families: P450 family (2), MFS transporters (2), PLAT/LH2 single domain proteins (4), and DNA-binding domain (DBD) lacking nuclear hormone receptor 96 (HR96) gene family (3), this region also included conserved secreted proteins (Csp) (2) and multiple chemosensory receptors from both the gustatory (TuGR) (1) and degenerin/epithelial Na+ channel (TuENaC) (6) families. Based on transcriptomic data of the genes within this region, three genes were constitutively upregulated after tomato selection: *tetur11g05520* (CYP385C4), *tetur11g05420* (Csp1e) and *tetur11g05740* (PLAT9); while five genes were constitutively downregulated: *tetur11g05540* (CYP385C3), *tetur11g05410*/*tetur11g05550* (MFS transporters), tetur11g05720 (PLAT10) and tetur11g05730 (PLAT11). As we suspect selection towards NL alleles (Supplementary Figure 6B), we filtered the SnpEff predictions (Supplementary Table 10) to assess the effect of variants originating specifically from the NL parental line. High impact variants were identified in *tetur11g05740* (PLAT9), *tetur11g05350* (TuENaC63) and *tetur11g05510* (TuGR332). Moderate impact variants, mostly missense variants, were observed in several genes, including *tetur11g05310* (TuENaC59), *tetur11g05330* (TuENaC61), *tetur11g05420* (Csp1e), *tetur11g05450* (Csp1f), *tetur11g05470* (HR96), *tetur11g05500* (HR96), *tetur11g05510* (TuGR332), *tetur11g05520* (CYP385C4), *tetur11g05540* (CYP385C3), *tetur11g05550* (MFS), *tetur11g5720* (PLAT10), *tetur11g05730* (PLAT11) and *tetur11g05740* (PLAT9). Notably, the largest differences in mean AF between tomato and bean replicates for these high- and moderate-impact variants were located in genes Csp1f and TuENaC63. Interestingly, a high-impact variant resulting in the gain of an early stop codon in TuENaC63 was selected in the tomato replicates.

##### 3.1.2.4 Transcriptomic changes upon tomato adaptation

To assess transcriptomic responses following tomato selection, we focused on two groups of DE genes (FDR < 0.05 and |log_2_FC| > 1). First, we detected genes that showed constitutive DE upon tomato adaptation. These are genes for which the selection host, bean or tomato, had a significant effect on gene expression in the populations feeding on bean (i.e., group 1). Second, we identified genes with significant differences in induced expression between bean and tomato replicates following the host shift from bean to tomato. These are genes for which the interaction effect between selection host and feeding host was significant (i.e., group 2).

In total, we found 190 genes in group 1 (Supplementary Table 11) and 12 genes in group 2 (Supplementary Table 12). Out of the 190 genes in group 1, the majority of the genes (144, 76%) were constitutively downregulated upon tomato adaptation. GO enrichment analyses on the set of downregulated genes revealed significant hits (FDR < 0.05) for the biological process proteolysis (GO:0006508), as well as the molecular functions cysteine-type peptidase activity (GO:0008234), heme-binding (GO:0020037), serine-type endopeptidase activity (GO:0004252), oxidoreductase activity (GO:0016705), iron-iron binding (GO:0005506) and steroid hormone receptor activity (GO:0003707). We checked whether the genes in group 1 showed significant DE upon exposure to tomato in the bean replicates (Supplementary Table 11). Out of the 190 genes in group 1, 139 did not exhibit plasticity in the bean-control populations.

All genes with a significant interaction term (group 2) had a higher expression in the tomato replicates compared to the bean replicates upon feeding on tomato. Interestingly, the MFS transporter (*tetur11g05410*), previously identified in section 3.1.2.3 as a candidate gene in the QTL2 region, ranked second highest in log_2_FC and lowest in p-value in the gene set of group 2. Additionally, the set included 2 lipocalin genes (ApoD11 and ApoD9), a CCE gene (TuCCE34), an UGT gene and 2 genes involved in digestion (TuPap-19 and SPH43).

The combination of genomic and transcriptomic data following adaptation allowed us to link genes with altered transcriptional responses to the detected QTL regions. Among the 155 constitutively DEGs (group 1) located on the 3 chromosomes (excluding those on loose scaffolds), 79 were found within significant QTL genomic windows. For each gene, we selected the windows overlapping its start position and calculated the average fraction of SNPs with significant differences (FDR < 0.01) in allele counts between tomato and bean replicates across all overlapping windows. Genes with an average fraction exceeding the genome-wide significance threshold across all overlapping windows were considered to be within the QTL regions. In total we found 76 DE genes (17 up, 59 down) located outside the QTL intervals, 79 (20 up, 59 down) were located inside the QTL regions. Out of the 12 genes showing a significant interaction effect (group 2), 6 were located within the QTL regions and 6 outside (Supplementary Table 12).

### 3.2 F2 backcross approach

We conducted a complementary approach to identify genomic regions linked to performance on tomato, using lines unrelated to the ones used in the experimental evolution approach: Pol-R and VR-BE, representing a low and high performer on tomato, respectively. We created a F2 backcross by crossing their F1 female offspring back to males of the low performer, Pol-R, and we contrasted pooled samples of high- and low-performing individuals with a pool of random individuals of the F2 backcross.

#### 3.2.1 Reproductive performance of parental lines and hybrids

The reproductive performance on bean did not differ between the parental lines and their F1 hybrid (∼4.5 eggs/female/day). On tomato, the VR-BE and its F1 hybrid laid approximately 6 eggs/female/day, and the Pol-R parent laid significantly fewer eggs, ∼2.5 eggs/female/day (Figure 5A; Supplementary Table 6). The reproductive output of the F2 cross on tomato was on average 4.24 ± 0.09 eggs/female/day (n = 933). However, the variation in egg number was spread across the large range for the F2 population, ranging from 0 to 12 eggs (Figure 5B). The distribution of egg numbers in the F2 population featured two peaks, around 2 and 6 eggs respectively, resembling the performance of each parental line. We pooled approximately 150 individuals from the F2 population belonging to the ∼15% of the distribution that laid the largest number of eggs (7-12 eggs/female/day), as well as approximately 150 individuals from the population that laid a low number of eggs, and 150 randomly selected individuals (control). We avoided collecting individuals that laid zero eggs to prevent inflating our genomic analyses with individuals with impaired physiology, and thus we pooled individuals that laid between 0.25 to 2.25 eggs/female/day (Figure 5B).

**Figure 5.**
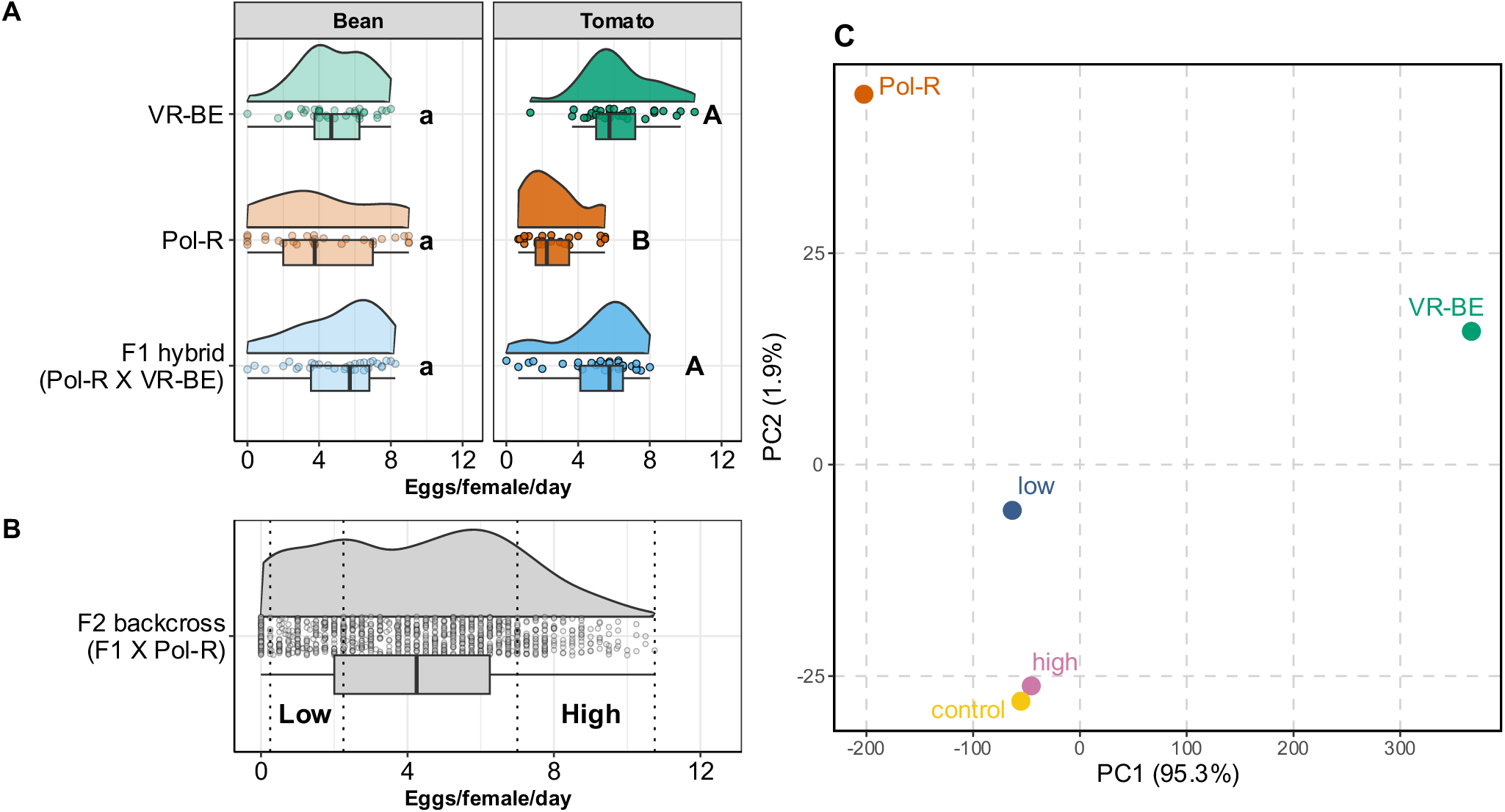
Phenotypic and genetic variation among parental lines and pooled samples in the F2 backcross approach. A) Reproductive performance on bean and tomato of females from two parental lines (VR-BE and Pol-R) and the F1 hybrid cross of Pol-R mothers vs. VR-BE fathers (n = 32). B) Reproductive performance on tomato of the females from the F2 backcross between F1 females and Pol-R fathers (n = 933). In A and B, the number of eggs laid per female per day alive (x-axis) is depicted as circles in the plots, with boxplots showing the median and quartiles of the data, and half-violin plots showing a smoothed distribution of the data. C) PCA using genome-wide allele frequency data at polymorphic sites. Each point represents a sample, labelled accordingly. The percentage of explained variance is given in parentheses for each principal component (PC1 on the x-axis, PC2 on the y-axis).

#### 3.2.2 Genomic responses linked to performance on tomato

A PCA using genome-wide allele frequency data from 659,784 SNP loci of both parental lines and the three pooled samples (low – high – control) showed a separation of the parental lines along the first dimension (PC1), which accounted for 95.3% of the variance (Figure 5C). The pooled samples clustered closer to the Pol-R parental line, as expected, given the experimental design in which the F1 hybrid was backcrossed to the Pol-R parent.

To detect genomic regions potentially involved in performance on tomato, we performed a Fisher exact test to compare the allele counts per locus between the high-performing and control pool, as well as between the low-performing and control pool. We calculated the fraction of significant SNPs (p-value < 0.01) in sliding windows. This revealed differences in the genomic profiles between the pools contrasting in performance and the control pool (Figure 6A). We focused on two significant peaks (referred to as QTLh for the high vs. control comparison and QTLl for the low vs. control comparison) to explore potential candidate genes associated with performance on tomato. These peaks were verified by an alternative BSA analysis (Supplementary Note 3; Supplementary Figure 7B).

**Figure 6.**
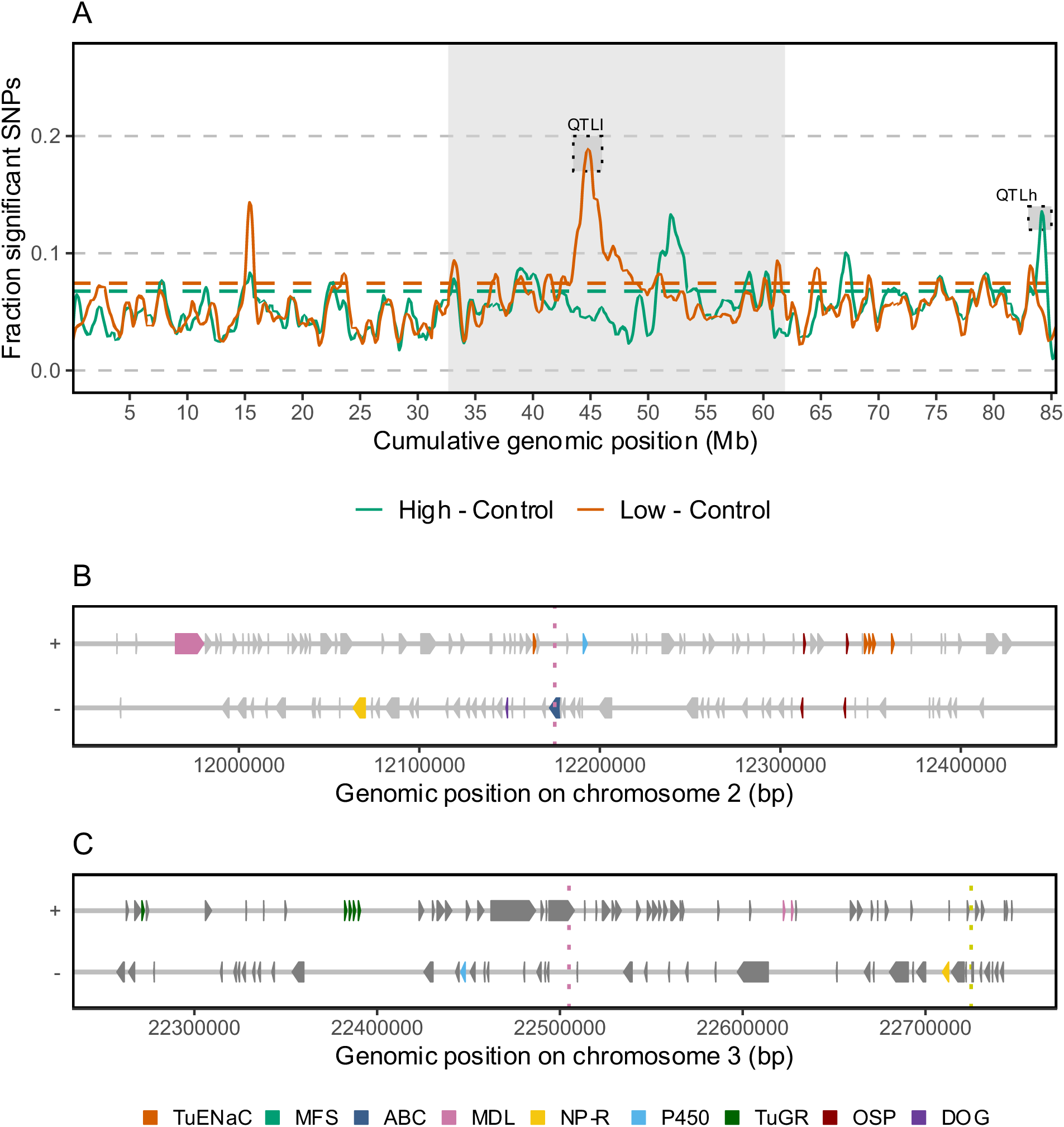
Contrast in genomic profile between high-performing and unselected (control) bulk and between low-performing and unselected (control) bulk. Panel (A) showing the fraction of significant SNPs (p-value < 0.01) calculated in sliding windows with window and step size set to 250kb and 25kb, respectively, for the high-performing vs control bulk (green) and the low-performing vs control bulk (red) The dashed red and green lines indicate the genome-wide significance threshold for windows as determined by random permutation of windows for the high-performing vs unselected, and the low-performing vs unselected bulk, respectively. Chromosomes are arranged in descending order and indicated by alternating shading. Genes within selected genomic regions around QTLl (B) and QTLh (C) are coloured (legend) according to their class if deemed interesting, otherwise they are shown in grey (TuENaC: degenerin/epithelial Na+ channel chemosensory receptors, MFS = major facilitator superfamily, ABC = ABC-transporters, MDL = MD-2-related lipid-recognition domain, NP-R: G-protein coupled neuropeptide receptor, P450 = cytochrome P450, TuGR = gustatory chemosensory receptor, OSP = Orphan secreted protein, DOG = intradiol ring cleavage dioxygenases). Vertical dotted lines represent the top genomic window in this experiment (blue) or the QTL peak detected in the same region in the Experimental Evolution approach (yellow).

The most prominent peak, QTLl on chromosome 2 at ∼12Mb (cumulative genomic position around 45 Mb) was associated with low reproductive performance. Analysis of AF differences of homozygous unique parental alleles revealed a decrease in VR-BE alleles in the low-performing pool compared to both the control and the high-performing pool (Supplementary Figure 8). We focused on a 500 kb region centred around the top window (Figure 6B, Supplementary Table 13), containing 123 genes. We identified genes within this region with high and moderate impact coding variants that showed higher AFs of the VR-BE allele in the high and control bulks compared to the low bulk (Supplementary Table 14). The variants with the largest differences in AF were located within a cluster of TuENaC genes (TuENaC13, TuENaC14, TuENaC15, TuENaC17), a complex, possibly copy number variable region, as suggested by DNA coverage. Another noteworthy candidate is *tetur01g00490* from the DOG family, previously shown to be highly upregulated in the VR-BE line when adapted to tomato compared to when adapted to bean (Njiru et al., 2023). We observed multiple missense variants within this gene and observed selection towards the Pol-R allele in the low-performing bulk, resulting in a loss of VR-BE alleles.

The QTLh region on chromosome 3, around 22-23 Mb (cumulative position ∼84 Mb) (Figure 6C, Supplementary Table 15), was associated with high reproductive performance. Within this rather complex region, with numerous moderate and high-impact variants (Supplementary Table 16), the most substantial AF differences were associated with a cluster of gustatory receptors (TuGR354, TuGR357, TuGR355, TuGR356). Overall, there was a trend of the alleles in the high-performing pool towards the VR-BE parent. Interestingly, this region was also associated with tomato selection in the Experimental Evolution approach (QTL3; Figure 4A). QTLh clearly overlapped with QTL3, with the top peaks of each QTL located only 220 kb apart (vs. ∼1 Mb peak widths). An analysis of the AF of unique parental marker SNPs in the Experimental Evolution approach indicates clear selection towards PL alleles in the tomato replicates within this region (Supplementary Figure 6C), along with the presence of missense variants with higher AFs in the tomato adapted populations.

### 3.3 Enrichment of gene families within QTL regions

In both experiments, chemosensory receptor genes (CR), TuGR and TuENaC, were the most abundant within the selected QTL regions. For the TuENac (p = 6.52e-08), but not for the TUGR (p = 0.15) family, this observation was supported by an exploratory hypergeometric statistical test for enrichment of these genes within the QTL regions.

## 4. Discussion

The fuel of adaptive evolution is the heritable variation upon which selection can act. Polyphagous herbivores harbour genetic and phenotypic variation related to host use within and between populations, which can be selected by the different potential hosts in their range. Here, we aimed to identify the molecular underpinnings of adaptation to, and performance on, a host plant in the extremely polyphagous mite *Tetranychus urticae* by exploring two complementary genetic mapping approaches. First, in an experimental evolution approach, we identified the molecular targets of host selection associated with adaptation in a genetically diverse mite population. Second, we conducted an independent F2 backcross experiment with parents that differ in host performance to map the genetic determinants of host performance. By comparing the results from both approaches, we were able to identify common genomic regions as well as those unique to the specific populations under investigation. The implications of our findings for understanding the evolution of adaptation in generalist herbivores are discussed.

### 4.1 Selection from tomato yields high performing individuals

The laboratory-created ‘Mix’ population had a high reproductive performance and fast juvenile development on both bean and tomato, surpassing most, but not all of its parents (Figure 2 and Supplementary Figure 3). Exposing the Mix to tomato varieties differing in their defence profiles revealed that female performance was similar on wildtype and JA-impaired tomatoes, suggesting that the Mix was resistant to JA-dependant induced tomato defences, even before experiencing tomato (Supplementary Figure 5B). This suggest an effect of heterosis, where hybridization between unrelated parents results in new combinations of beneficial alleles and purging of deleterious ones in the hybrid population (Martin-Roy et al., 2021).

Experimental evolution on tomato resulted in females with higher reproductive output and faster development than mites kept on bean, creating a pattern of local adaptation without apparent costs to tomato adaptation, as performance was high on both hosts for tomato-adapted females (Figure 2; Gould, 1979; Kawecki & Ebert, 2004; Magalhães et al., 2009; Wybouw et al., 2015). When tomato mites were reverted to bean for one or two generations, oviposition on tomato was lower than non-reverted mites, but still higher than bean mites on tomato. This suggests genetic evolution as the cause of increased oviposition, confirmed by our genomic analyses (see section 4.2).

Interestingly, we found that re-acclimatization to bean during juvenile development following tomato selection resulted in the highest adult reproductive performance, since mites reverted to bean for one generation showed the largest number of eggs on bean (Figure 2C). However, performance decreased after two generations (Figures 2D), suggesting a cost to maintaining high oviposition in an environment that does not select for it. A similar pattern was observed for juvenile survival, where mites reverted to bean for two generations had quicker development on bean and tomato, but lower survival on bean, and not on tomato, than any other treatments (Supplementary Figure 4). Whether maintaining a high performance is costly in a relatively ‘easy’ environment after selection on a challenging one, remains a possibility to be explored. Our results show that these costs may be spread between different developmental stages of the adapted organism, and thus that loci regulating juvenile development and adult performance are probably uncoupled and free to evolve independently from each other, as observed in natural populations (Villacis-Perez et al., 2021).

Mites kept on bean had a similar performance to when the Mix was first created (Figure 2; Supplementary Figure 3), suggesting that selection had reached a peak on the fitness landscape of bean. Tomato-adapted mites performed better than bean mites on tomato, bean and maize, but not on honeysuckle or on pepper, despite it being phylogenetically closer to tomato (Supplementary Figure 5A). Transcriptomic responses of *T. urticae* differ when feeding from tomato or maize, but the expression of multigene detoxification families is less host-specific than individual genes, thus suggesting a commonality in detoxification pathways between hosts (Snoeck et al., 2018). Our results do not exclude the possibility that mechanisms related to nutrition acquisition evolved upon tomato selection. Together, these observations suggest that tomato selects for individuals with high performance across hosts beyond tomato, while bean relaxes selection, allowing populations to harbour individuals with slower development and lower performance. In nature, challenging hosts may act as filters that yield populations with a higher performance and thus with the potential of becoming more perjurious pests. High host performance is common in natural mite populations (Kant et al., 2008; Villacis-Perez et al., 2021). In the F2 backcross approach, we found evidence for the dominant inheritance of performance on tomato (Figure 5B). Natural variation in the expression of genes involved in allelochemical detoxification is often *trans*-regulated and inherited as a dominant trait in this species, and thus is open for selection when it is exposed in heterozygous diploid females (Ji et al., 2023; Kurlovs et al., 2022).

### 4.2 Molecular determinants of mite adaptation

While several pesticide resistance mechanisms have been readily mapped in *T. urticae* (De Beer et al., 2022; Fotoukkiaii et al., 2021; Vandenhole et al., 2023; Villacis-Perez et al., 2023; Wybouw et al., 2019), the alleged polygenic nature of host plant adaptation in this species has proven harder to dissect. In Wybouw et al., 2019, a BSA on a population from a biparental cross of inbred lines identified a broad region on chromosome 3 of *T. urticae* that responded to tomato selection. Interestingly, the two major QTL found in this study, QTL1 and QTL2 (Figure 4), overlap with this region on chromosome 3. In our study, we detected multiple distinct peaks within this region, allowing us to propose a list of candidate genes for functional validation.

Following experimental evolution, we observed larger variation between the genomic profiles of tomato replicates compared to the bean replicates (Figure 3A). This could be an intrinsic property of the selection regime imposed by tomato, where bottlenecks and subsequent genetic drift mark differences among the experimental replicates, particularly since the fitness of the Mix population was relatively lower on tomato than on bean upon its creation. In contrast, the bean populations likely did not experience such bottlenecks, as bean was not a challenging host for the Mix, which developed and expanded on bean. Supporting this idea, we found that the average genome-wide heterozygosity levels within replicates were significantly higher in the bean compared to tomato replicates.

Among the candidate genes responding to tomato selection was CYP392B3, located at the peak of QTL1 (Figure 4B). This gene was previously reported to be significantly upregulated in multiple *T. urticae* populations feeding on tomato (e.g., log2FC of 4.78 in Snoeck et al., 2018; log2FC of 3.46 in Wybouw et al., 2015). Notably, Njiru et al., 2023 propagated the VR-BE line (used here for the F2 backcross approach) on tomato and on bean, and found that the tomato line had a higher expression of CYP392B3 (log2FC of 4.2). In our study, this gene did not show constitutive upregulation or altered induced expression upon tomato adaptation. However, we identified an additional copy of the CYP392B3 gene in the genome of the parental line PL, which was selected in the tomato replicates, suggesting that variation in its coding sequence could be the target of tomato selection.

Chemosensory receptor genes (CR), including TuGR and TuENaC, stood out as prominent candidates in both experimental approaches. The TuENaC family showed significant enrichment within all QTL regions across chromosomes 1, 2, and 3, relative to all genes from this family across the genome. These receptor genes have been implicated in host plant adaptation not only because generalist arthropod species often have larger repertoires of CR genes than specialists (L. Chen et al., 2023; R. Chen et al., 2024; Ngoc et al., 2016; Xu et al., 2016), but also due to extensive changes and pseudogenization of CR genes linked to host plant speciation (Anholt, 2020; Goldman-Huertas et al., 2015). More specifically, for generalists, sensory changes have been suggested to involve a loss of function rather than a gain of function (Cohen et al., 2024). Strikingly, we found that tomato selected for a stop codon gain in TuENaC63 within QTL2. This resembles the patterns associated with the rapid evolutionary changes in the chemoreceptor family of *Drosophila sechellia*, which prefers to feed on *Morinda citrifolia*, a plant avoided by other *Drosophila* species (Auer et al., 2020). A premature stop codon in the Obp56e gene, producing a loss-of-function allele, influenced host preference behaviour in this species, while RNAi-mediated knockdown of Obp56e in *Drosophila melanogaster* resulted in reduced avoidance of *Morinda citrifolia* (Dworkin & Jones, 2009). We propose TuENaC63 in *T. urticae* as a candidate for functional validation, which can be efficiently performed in this species using CRISPR/Cas9 (De Rouck et al., 2024).

Detoxification genes are often induced by arthropods upon host shifts, but whether these genes drive the evolution of host adaptation is unclear (Rioja et al., 2017). Two main hypotheses explain how induced gene expression can promote adaptation. The first hypothesis assumes that gene expression is independent of the environment, with adaptation occurring primarily through genetic changes in constitutive expression. In this view, genes that do not show initial transcriptional plasticity are impacted by selection equally as those that do. The second hypothesis suggests that exposure to a new environment induces a gene expression response, and adaptation acts on this through genetic assimilation and genetic accommodation. In both processes, selection primarily acts on genes that show initial transcriptional plasticity. Genetic assimilation involves the induced expression response becoming constitutively up- or downregulated. Genetic accommodation involves changes in the degree and pattern of plasticity (P. Chen & Zhang, 2023; Ehrenreich & Pfennig, 2016; Wybouw et al., 2015). Following experimental evolution, we identified 51 genes associated with genetic assimilation (Supplementary Table 11; constitutively DE genes upon tomato adaptation which also showed DE upon tomato exposure in bean replicates) and 12 genes associated with genetic accommodation (Supplementary Table 12). The majority of genes with constitutively altered transcriptional profiles upon adaptation (139 genes) did not show an initial response to tomato, confirming previous findings by Wybouw et al., 2015. A similar pattern was observed in a study on transcriptional responses of the generalist *Bemisia tabaci* adapted to habanero pepper, where most changes were categorized as evolved rather than plastic responses (Tadmor et al., 2022). Similar to the findings of Wybouw et al., 2015, we found that most genes that were constitutively DE upon tomato adaptation (group 1) were downregulated (76%). However, unlike this previous study, we identified enriched processes related to digestion and detoxification within this set of downregulated genes. Together with the observation that taste perception genes are selected upon tomato adaptation, our data suggest that concerted changes in the identity and expression of genes involved in detoxification, nutrient acquisition and taste perception are important factors for host use and adaptation.

Integrating genomic and transcriptomic data allowed us to identify differentially expressed genes inside and outside genomic regions under selection, which provides further cues on the role of *cis*- and *trans*-regulation in adaptive expression evolution. Genes outside selected regions are likely *trans*-regulated by diffusible elements such as transcription factors, in contrast to genes within QTL regions, which are potentially influenced by selection on *cis*-regulatory elements or copy number variation (Signor & Nuzhdin, 2018), although regulation by a nearby encoded *trans* factor can still be possible. However, both *cis* and *trans* regulated DE genes may be due to expression quantitative trait loci (eQTLs) linked to causal loci under selection. Our study found a relatively balanced distribution of putatively *cis*-regulated (79 genes) and *trans*-regulated (76 genes) genes. This contrasts with Fotoukkiaii et al., 2021 who reported that only 31% of DE genes between pyflubumide-selected and control *T. urticae* populations were within QTL genomic intervals and thus putatively *cis*-regulated. Additionally, it differs from the findings of Kurlovs et al., 2022, who suggested abundant *trans* regulation for *T. urticae*. The results of Kurlovs et al., 2022 align with the hypothesis that *cis* effects predominate in interspecific comparisons, while *trans* effects are more common in intraspecific comparisons, and thus are expected to play a larger role in expression variation in the context of host plant adaptation (Benowitz et al., 2020; Signor & Nuzhdin, 2018). The diverse genetic background used in this study is more complex compared to the biparental inbred line populations used in other studies, potentially leading to more epistatic effects and compensatory *cis* and *trans* effects, which in turn might mask *trans* effects.

## 5. Conclusion

In nature, populations of *Tetranychus urticae* form mosaics of genetic and phenotypic variation. Here, we address the question whether different plant species select for genetic variants available in the genetic pool of this generalist herbivore, and whether we can capture these genetic variants using different empirical approaches. This complementary approach was useful to dissect between the evolved genetic changes due to experimental host selection and the genetic elements associated with host performance that segregate in natural populations. Our experimental evolution pipeline was able to dissect regions under selection to levels previously not achieved, presenting a list of candidate genes to be functionally validated to understand their role in performance across host plant species. Populations experimentally adapted to tomato evolved higher reproductive performance following selection, and strikingly, this fitness advantage was not restricted to the host of selection, tomato, but extended to other hosts as well. Furthermore, our study highlights the diversity in mechanisms associated with host adaptation across populations of this generalist species, which indicates that there is no single genetic architecture underlying host plant adaptation, but rather, a diversity of mechanisms available for this species to colonise novel plants.

## Author contributions

TVL and EVP designed the experiments. EVP, BDB, SAA, and RdJ conducted the experiments and analysed the data together with FDG with contributions from TDM. EVP and FDG wrote the manuscript with contributions from TDM and TVL. All authors read and approved the final manuscript.

## Acknowledgments

This work was supported by the Research Council (ERC) under the European Union’s Horizon 2020 research and innovation program [ERC grant 772026-POLYADAPT to TVL] and the Research Foundation Flanders (FWO) [grant 3G006720 to TDM, grant G017923N to TVL].

## Data availability statement

WGS and RNA-seq raw data will be deposited in appropriate repositories upon acceptance.

## Benefit-sharing statement

International collaboration has been fundamental to the successful completion of this work and has contributed to sharing of scientific results, cooperation, education and training of early career researchers across countries. In addition, the research addresses a priority concern for the public, in this case the adaptive potential of a widespread pest. Benefits from this research include the availability of data and results on public databases, as described above.

