## Supplementary material for "Independent genetic mapping experiments identify diverse molecular determinants of host adaptation in a generalist herbivore"

* Shared first authorship

### Supplementary Note 1

#### DNA extraction and whole-genome sequencing

DNA was obtained from pooled samples by a chloroform-phenol extraction as described previously (Villacis-Perez et al., 2021). The quality and amount of DNA per sample were first assessed using a Implen NanoPhotometer N60 (Implen, Germany) and by visual assessment of gel electrophoresis (1% agarose gel; 45min; 50V).

Sequencing libraries for the parental lines and the Mix were constructed using the NEBNext Ultra II Library Prep Kit (PCR-free) for Illumina at the NXTGNT sequencing facility (Ghent University, Ghent, Belgium), followed by sequencing at Genewiz (Leipzig, Germany) on an Illumina Hiseq3000 platform for the parental lines, and sequencing at NXTGNT (Ghent University, Ghent, Belgium) on an Illumina Nextseq500 platform for the Mix. For the five selected replicates from the Mix BSA, libraries were constructed using the TruSeq Nano DNA Kit (Illumina, USA) and sequenced on an Illumina NovaSeq6000 platform at Macrogen (Amsterdam, The Netherlands). Samples for the F2 approach were sent to Novogene (Cambridge, UK) for library preparation using the Novogene NGS DNA Library Prep Set and subsequent sequencing on an Illumina NovaSeqX platform. Each of the above-mentioned sequencing runs yielded 150 bp paired-end reads, with a minimum of 25 million reads per sample, except for the sequencing run of the Mix, which yielded 110 million 75 bp paired-end reads.

#### Genome alignment, variant calling and PCA

All DNA reads were aligned to the London reference genome (Wybouw et al., 2019) using BWA (0.7.17-r1188) (Heng Li, 2013) with default settings. The resulting bam files were sorted by coordinates using SAMtools (v1.6) (Danecek et al., 2021), and duplicates were marked with Picard (v2.25.0) (https://broadinstintute.github.io/picard). Genetic variants were jointly called across all samples of each respective approach, following GATK's (v4.2.0.0) best practices workflow (McKenna et al., 2010), and subsequently filtered according to GATK's recommended criteria for hard filtering of germline variants: 1) MQ ≥ 40, 2) QD ≥ 2, 3) Q ≥ 100, 4) SOR ≤ 3, 5) FS ≤ 60, 6) MQRankSum ≥ -12.5, 7) ReadPosRankSum ≥ -8. Additionally, only bi-allelic SNPs having a minor allele frequency (MAF) across all samples ≥ 0.1 and read coverages at variant sites within 25-150% of the respective sample's genome-wide mean, were retained. For these loci, the allele frequency (AF) of the alternate allele per locus was calculated for each sample. To assess the genetic variability between and within the samples, these genome-wide AFs were used for a principal component analysis (PCA) using the Python (v3.10.6) package 'scipy' (v1.11.4) (Virtanen et al., 2020). The average genome-wide heterozygosity percentage for each line was determined using a sliding window approach (window size = 100kb, step size = 10 kb), where the number of heterozygous SNPs in each window were calculated and these values were then averaged over all genome-wide windows.

#### RNA sequencing, alignment and differential gene expression analysis

RNA was extracted using the RNeasy Plus mini kit (Qiagen, Germany) following the manufacturer's protocol. Quantity and quality of the extracted RNA was assessed using a DeNovix DS-11 spectrophotometer (DeNovix, USA) and by visual inspection of gel electrophoresis (1% agarose gel; 45 min; 50 V). The sequencing libraries were constructed using the Illumina stranded mRNA ligation kit (Illumina, USA) at Novogene (Cambridge, UK), and subsequently sequenced on an Illumina NovaSeqX platform to generate 150 bp paired-end reads. The RNAseq reads were aligned to the London reference genome using STAR (v2.7.9a) (Dobin et al., 2013) in two-pass alignment mode, with a maximum intron size of 30 kb, and using the GFF annotation provided by (Ji et al., 2023). The resulting BAM files, sorted by chromosomal coordinates, were subjected to gene-level read counting with HTSeq (v2.0.2) (Anders et al., 2015), accounting for strand-specificity and using the above-mentioned annotation. These counts were filtered using the default settings of the *filterByExpr* function in the R (v4.3.1) package 'edgeR' (v3.42.4) (Robinson et al., 2010), and normalized using the trimmed mean of M-values (TMM) algorithm, prior to differential expression (DE) analysis using edgeR's quasi-likelihood F-tests. PCA was performed on the log-transformed normalized counts of the 500 most variable genes across all samples using the function *prcomp* from the R package 'stats' (v4.3.1). A blocking factor for the replicate ID was included in the model due to the paired experimental design. Differentially expressed genes (DEGs) were selected based on an absolute log2 fold change >1 and a Benjamini-Hochberg (BH) adjusted p-value < 0.05.

#### Gene ontology (GO) enrichment analysis

GO enrichment analyses were performed with the *enricher* function of the package 'clusterProfiler' in R (v4.8.3), using a BH-adjusted p-value cut-off of 0.05. The background gene set consisted of the remaining genes after applying the edgeR *filterByExpr* function, thus excluding low-count genes, which were also not included in the DE analysis. The Molecular Function (MF) and Biological Process (BP) terms were sourced from the ORCAE database (v01252019) (Sterck et al., 2012).

#### Effect of variant alleles in coding sequences

To evaluate the potential impact of variant alleles on loci identified in the genomic scans, we predicted the coding effects of SNPs and small indels detected by the GATK variant calling on the parental lines using SnpEff (v5.1f) (Cingolani et al., 2012). We used the *T. urticae* coding sequence database available in the SnpEff pre-built databases, which is based on the June 23, 2016 annotation available from the Online Resource for Community Annotation of Eukaryotes (ORCAE) (Sterck et al., 2012). Custom Python (v3.10.6) scripts were subsequently used to verify the presence of high- and moderate-impact variants in the parental lines.

#### Enrichment of gene families in QTL regions

For interesting gene families, we tested whether the gene family was significantly enriched within the selected QTL regions by performing a hypergeometric statistical test using the R function *dhyper*. Essentially, we identified the total number of genes genome-wide that belonged to a particular family. We then sampled genes from the QTL regions and identified how many of these belong to the specific family. The hypergeometric test determined whether the observed number of family-specific genes within the QTL regions was significantly higher than what we would expect by chance, given the total number of such genes in the genome.

### Supplementary Note 2: Phenotyping of fitness traits in the Experimental Evolution approach

#### Phenotyping fitness traits of parental lines and the Mix

To infer the extent of variation in traits associated with the use (and adaptation) to different host plants in the parental lines and the Mix, adult female reproductive output, juvenile survival, and juvenile development until adulthood were measured. To synchronise the age of experimental individuals, 100-150 adult, mated females from each parental line and from the Mix population were collected with a vacuum pump and transferred to detached bean leaves. Two days after, adult females were removed from the leaves. After twelve days, females and males had reached adulthood and mated on the leaves. Thus, experimental females were 2 ± 1 days old when used for the experiments.

To assess the differences in the reproductive output of gravid females across lines and across plants, age-synchronised females were each transferred to a 15mm diameter leaf disc of either bean or tomato placed on wet cotton wool (n = 32 females per line per plant). Survival was scored daily for a total of 4 days. Each female was scored with a 1 if alive, with a 0.5 if she was stuck to the wet cotton (in which case the female was relocated back into the centre of the disc), or 0 if the female was dead. After 4 days, the total number of eggs per disc was counted, and the total number of eggs was corrected for mortality. To analyse the data, first an omnibus analysis was conducted fitting a linear model to the parental data, with an interaction between line and plant species. Then, the number of eggs either between plants and within each parental line, or within plant and between all the parental lines were compared. The Mix was analysed separately, and thus it was only compared within line and between plants. Independent linear models were fitted to these datasets. All analyses were conducted using the package 'aov' in R (v4.3.2) and the fit of each model was assessed visually by plotting its residuals and fitted values. The data were transformed whenever model assumptions of residual normality and homoscedasticity were not met.

To assess differences in the probability of survival and development until adulthood of juvenile individuals, arenas of either bean or tomato of 24mm in diameter were created and six adult, mated females of a synchronised cohort were transferred to each disc, for a total of six discs per line, per plant. Twenty four hours later, all eggs and web were removed from the discs. After another 24 hours, the females were removed, and the total number of eggs was reduced to 12 eggs per disc to facilitate tracking of each individual, for a total of 72 juveniles per line, per plant. The survival and developmental stage of each individual was scored at five points, starting at day 5 after removing the adult females and thereafter every two days up to day 13, which coincides with the time needed for egg hatching and with the expected developmental time to adulthood of this species under standard conditions, respectively (Hodek, 1987). For the development to adulthood, the focus was on the individuals alive per time point. Both the probability of survival and of reaching adulthood were compared either between plants and within each line, or between lines within each plant. To analyse the statistical differences in juvenile survival and development, a Cox proportional hazards model was fitted to the censored data and assessed with a long-rank test (package 'survival' v3.5-7) in R (v4.3.2), with either line or plant as the main factor.

#### Phenotyping fitness traits after experimental evolution

To discern between genetic evolution and acclimatisation or potential maternal effects, we defined four treatments: 1) mites from bean; 2) mites from tomato; 3) tomato mites reverted for one generation to bean; 4) tomato mites reverted for two generations to bean. To minimise experimental variation in the quantification of the fitness traits, such as due to variation in plant quality during the phenotyping period or due to other unforeseen factors, mite cohorts were synchronised so that treatments 1, 2 and 3 were conducted simultaneously for each replicate. Individuals from treatment 4 (tomato mites reverted to bean for two generations) were conducted one generation after the batch from treatments 1-3. For example, ~150 females from replicate 1 on tomato were transferred to a detached bean leaf and ~150 to a tomato leaf, and ~150 females from its paired bean replicate 1 were transferred to a bean leaf, in parallel. Females were allowed to lay eggs for 2-3 days and then were removed. Fourteen days later, adult females of the synchronised cohorts were transferred to experimental arenas of bean and tomato, as the ones described in section 2.3. At the same time, ~150 females from the tomato replicate that had grown on bean were collected and transferred to a new bean leaf. After 2-3 days of egg laying, females were removed from their leaves. Fourteen days later, individuals from the tomato replicate that were reverted to bean for two generations (treatment 4) were transferred to experimental arenas as the ones described in section 2.3. Each replicate batch was offset by one day, so that e.g., replicate 2 started one day after replicate 1, and so forth until reaching replicate 10. The data was analysed and checked for model fit as specified in section 2.3, except that for assessing the reproductive output of females, where a linear mixed model was fitted to the dataset using the command *lmer* within package 'lme4' v1.1.35-1 in R 4.3.2, with either treatment (levels 1-4) as main factor and replicate (1-10) as a random factor for comparisons across treatments within a plant, or with plant as the main factor and replicate (1-10) as a random factor for comparisons within each treatment and between plants.

#### Phenotyping reproductive performance on alternative host plants after experimental evolution

After approximately 30 generations of experimental evolution, a bioassay was conducted using a subset of three paired populations (replicates 6, 7 and 9). We synchronised the age of mites and replicated three of the four previous treatments: mites from bean, mites from tomato, and mites from tomato reverted to bean for two generations. Female reproductive output was scored on leaf discs of bean, tomato, maize, honeysuckle, and sweet pepper using the same set-up as described above (n = 32 per replicate, per treatment, per plant). A linear mixed model was fitted to the data using the command *lmer* within package 'lme4' v1.1.35-1 in R v4.3.2, with treatment (levels 1,2, 4) as main factor and replicate (6, 7, 9) as random factor. In a separate experiment, the reproductive output of age-synchronised females from replicate 7 was scored, in three treatments created with detached tomato leaves with the petiole dipped in a tube of a solution. Treatments included wildtype tomato (cv. Castlemart) dipped in water, a tomato mutant with impaired jasmonate production (cv. Def-1) dipped in water, and the same tomato mutant but dipped in a solution of jasmonic acid + isoleucine to rescue jasmonate production, following Ataide et al., 2016. Individual mites were isolated to each leaflet of a tomato leaf using wet tissue barriers, and a total of 5 females per leaf, and 6 leaves per treatment were used for the experiment. A linear mixed model was fitted to the data using the command *lmer* within package 'lme4' v1.1.35-1 in R v4.3.2, with treatment as main factor and leaf as random factor.

### Supplementary Note 3

#### Empirical FDR estimation GLM

Permuted datasets were generated to establish an empirical null distribution, reflecting the same level of overdispersion as the observed data, essentially reflecting a drift-only model as described by Hoedjes et al., 2019. Ten permuted datasets were created by randomizing the labels of the count dataset with certain restrictions: samples could not retain their original labels and original replicate pairs could not be pairs again in the permuted datasets. For each permuted dataset, p-values were computed using the same binomial GLM model and LRT. Subsequently, for each observed p-value and each permuted dataset individually, the number of permuted p-values lower than the observed value was calculated. Averaging across these ten calculated false discovery rates (FDR) yielded a mean FDR for each SNP. SNPs were considered significant if the mean FDR was below 0.01.

#### Standard and modified BSA script

The "RUN_BSA1.02.py" script (available at https://github.com/rmclarklab/BSA) was modified to assess AF differences in genomic windows with window and step size set to 150kb and 15kb (Experimental Evolution approach) or 250kb and 25kb (F2 backcross approach), respectively. The original script was designed for a biparental (inbred) cross, and the permutation approach used to assess significant QTLs from a genome-wide threshold of AF differences needed to be adjusted to run without providing parental information, and thus using all alleles, independent of the parental origin. To establish a genome-wide threshold, 10,000 permutations were performed using a false discovery rate (FDR) of 0.05.

We chose to use the method described in the main methods section 2.2.2 as we believe it has higher power in detecting QTLs, likely due to the conservative nature of the permutation approach in BSA method to distinguish peaks resulting from selection from those resulting from drift. Particularly when the baseline difference in allele frequency between bulks is higher, as expected in cases with multiple loci exhibiting (smaller) effects due to a polygenic architecture, the applied permutation approach might be too stringent. Additionally, if the starting allele frequency in the population of the selected allele was low, resulting in subtle differences in allele frequency after 20 generations, these limited differences may be overlooked by the standard BSA method.

### Supplementary figures

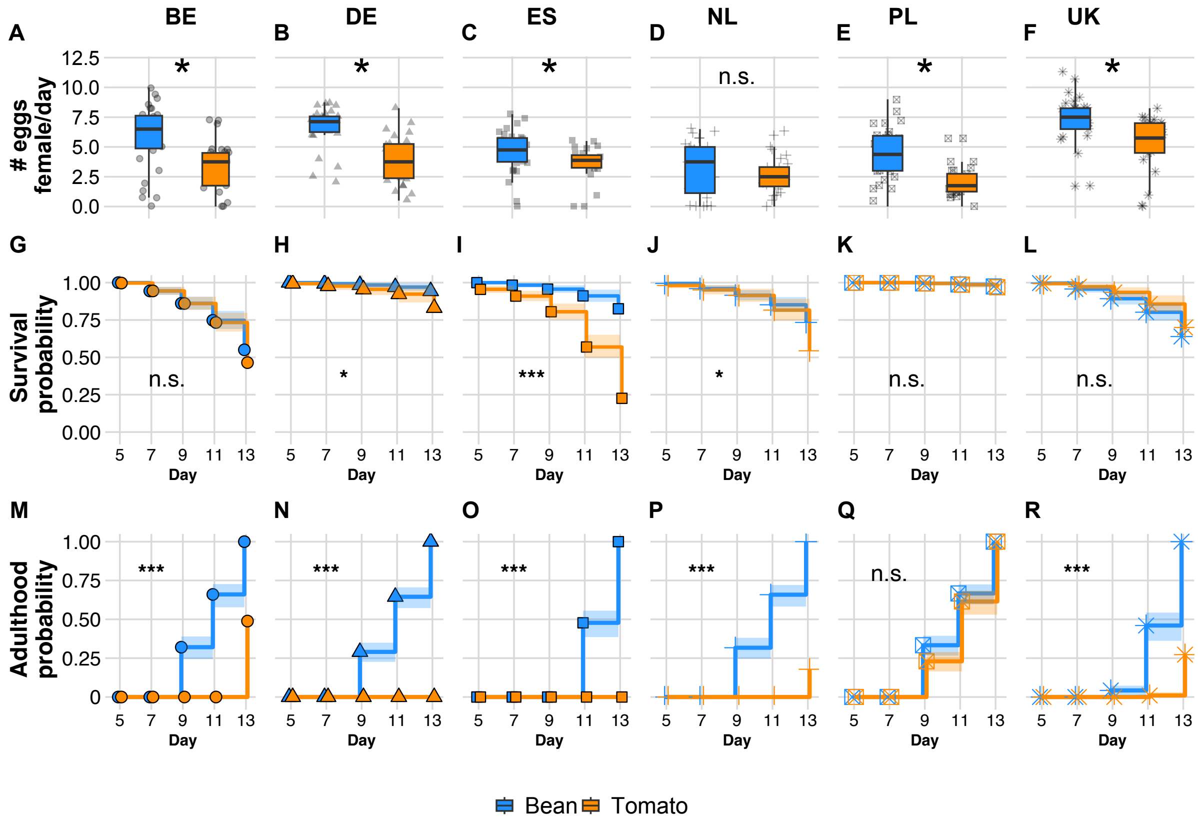


**Supplementary Figure 1. Phenotypic variation in fitness traits related to host use within parental lines.** Reproductive output of adult females (A-F), survival (G-L) and developmental time to adulthood of juveniles (M-R) of the six single-female expansion parental lines, on bean and tomato (legend). Reproductive output (A-F) is presented as the number of eggs laid per individual female per day alive (y-axes) per plant (x-axes, legend), with boxplots showing the individual data points (grey shapes), the median (black line within the interquartile box), and the bottom and top 25% quartiles of the data values (whiskers). Stars show a significant difference in the reproductive output between plants, according to a linear model fitted to the data within panel. Survival (G-L) and developmental time to adulthood of juveniles (M-R) is presented as the probability (y-axes) of surviving or reaching adulthood, respectively, across five time points spanning from egg hatching until adulthood, per plant (legend). Stars show a significant difference between plants in the probability of a line's survival or development, according to a survival analysis based on a Cox proportional hazards model per panel.

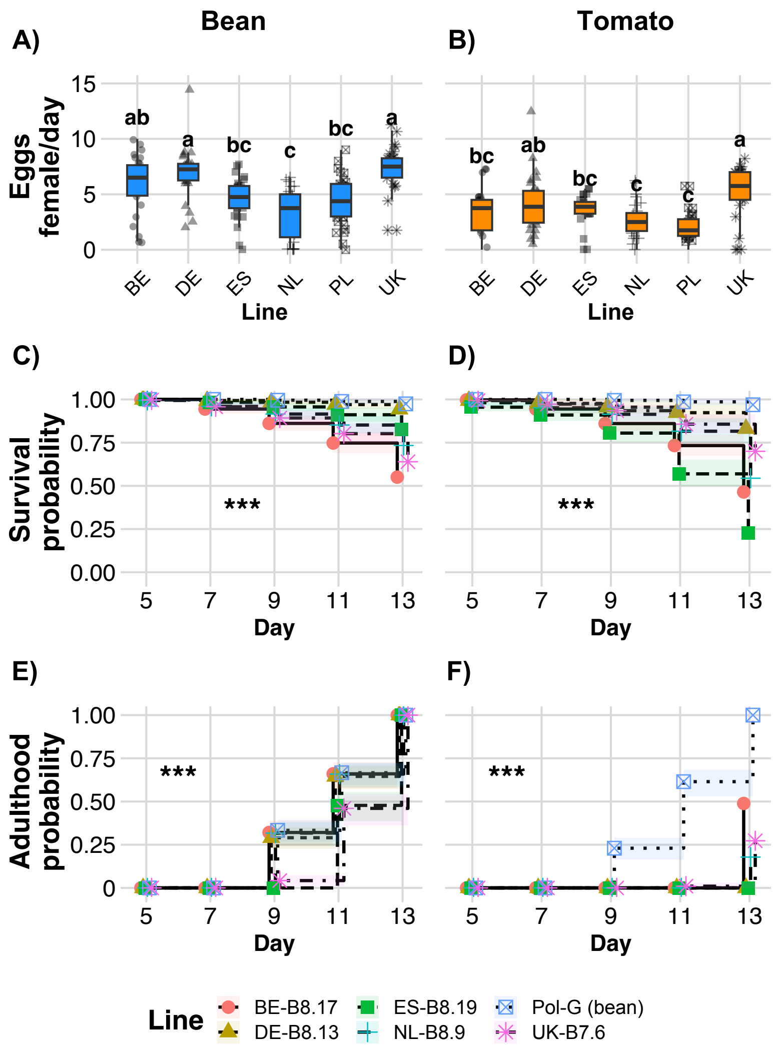


**Supplementary Figure 2. Phenotypic variation in fitness traits associated to host use among six lines of *Tetranychus urticae*.** Reproductive output of adult females (A-B), survival (C-D) and developmental time to adulthood of juveniles (E-F) compared across the six single-female expansion parental lines (legend), on bean (A, C, E) or tomato (B, D, F). Reproductive output (A-B) is presented as the number of eggs laid per individual female per day alive (y-axes) per line (x-axes) with boxplots showing the individual data points (grey shapes), the median (black line within the interquartile box), and the bottom and top 25% quartiles of the data values (whiskers). Letters on top of each boxplot show a significant difference in the reproductive output between lines, according to a linear model fitted to the data within panel. Survival (C-D) and developmental time to adulthood of juveniles (E-F) is presented as the probability (y-axes) of surviving or reaching adulthood, respectively, across five time points spanning from egg hatching until adulthood, per line (legend). Stars show a significant difference between lines in the probability of survival or development, according to a survival analysis based on a Cox

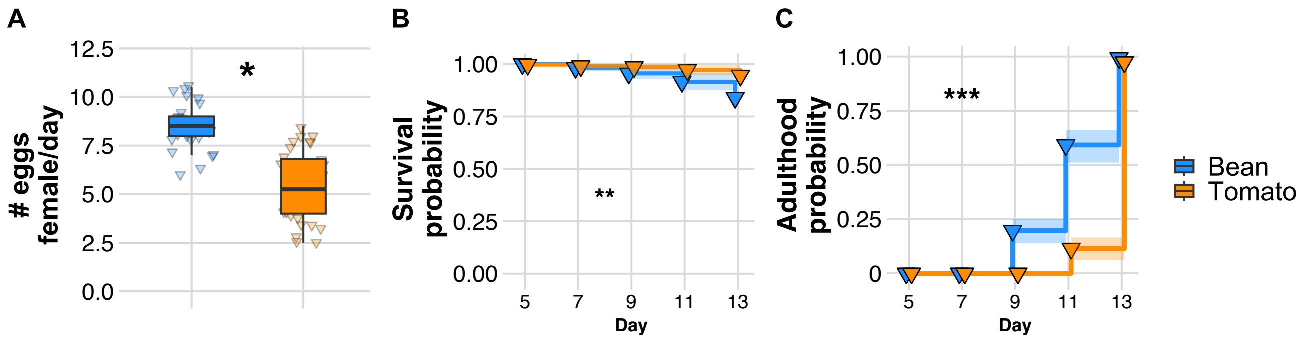


**Supplementary Figure 3. Fitness traits of the Mix population on bean and tomato.** Female reproductive output (A) is presented as the number of eggs laid per individual female per day alive (y-axis) per plant (x-axis, legend), with boxplots showing the individual data points (shapes), the median (black line within the interquartile box), and the bottom and top 25% quartiles of the data values (whiskers). The star shows a significant difference in the reproductive output between plants, according to a linear model fitted to the data. Survival (B) and developmental time to adulthood of juveniles (C) is presented as the probability (y-axes) of surviving or reaching adulthood, respectively, across five time points spanning from egg hatching until adulthood, per plant (legend). Stars show a significant difference between plants in the probability of the line's survival or development to adulthood, according to a survival analysis based on a Cox proportional hazards model fitted per panel.

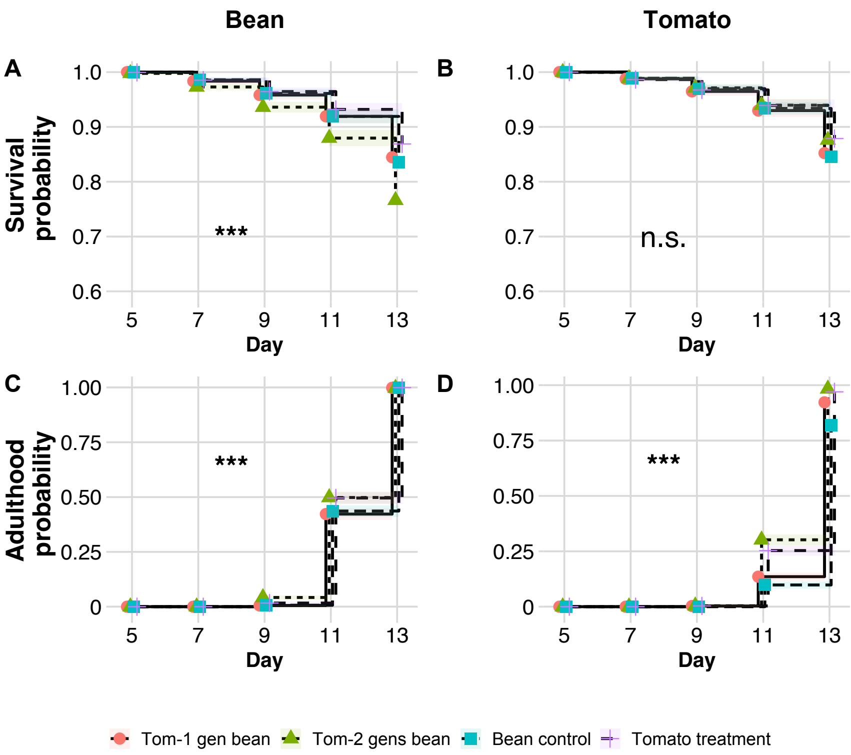


**Supplementary Figure 4. Survival and development to adulthood of juveniles after 20 generations of experimental evolution.** The probability (y-axes) of survival (A-B), and the probability of reaching adulthood (C-D) of juveniles on bean (A-C) and tomato (B-D) from the unselected control bean lines, the tomato-evolved lines, and the tomato-evolved lines after being reverted to bean for either one or two generations (legend), across five time points (x-axes) spanning from egg hatching at day 5 until day 13. Stars within each panel show a significant difference in the probability of survival or development of each of the mite lines according to a survival analysis based on a Cox proportional hazards model fitted per panel.

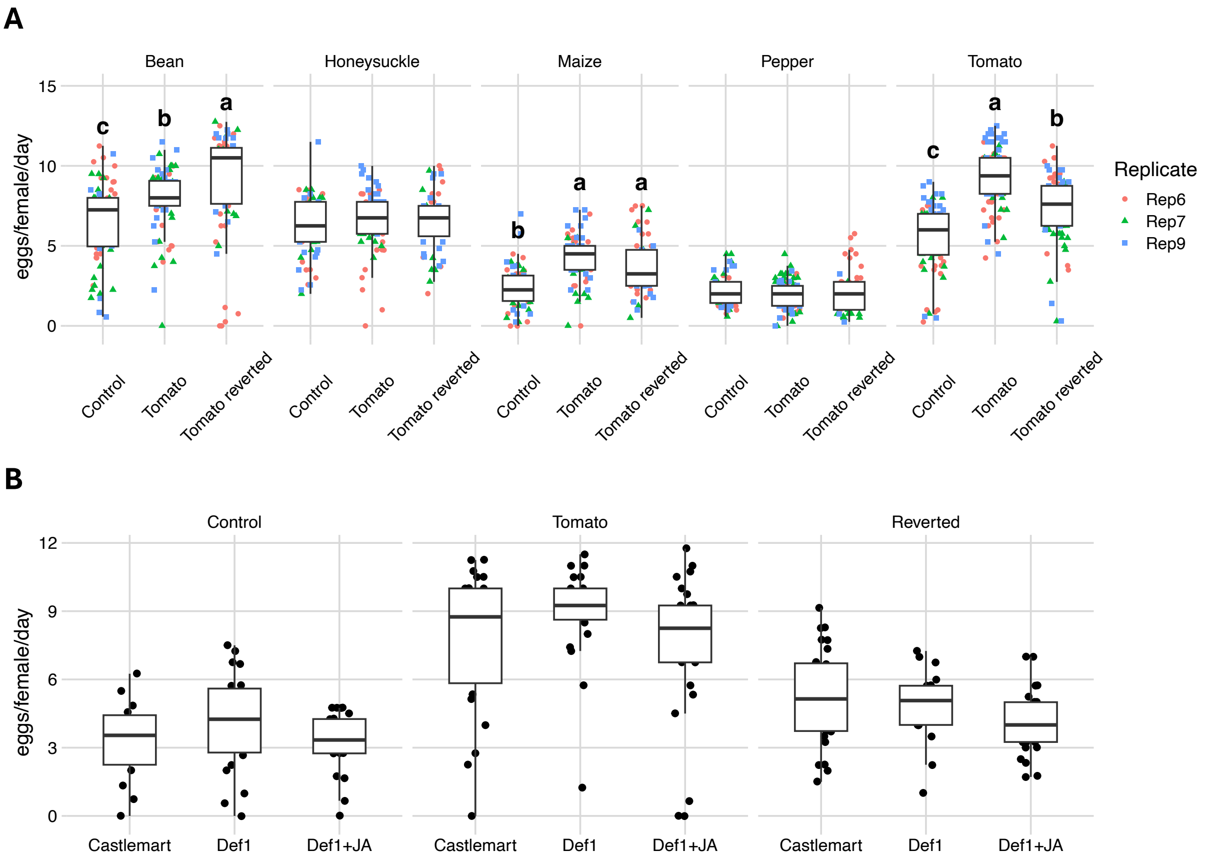


**Supplementary Figure 5. Reproductive performance of adult females after 30 generations of experimental evolution.** A) Reproductive performance of females after 20 generations on tomato, on plants of bean, honeysuckle, maize, pepper and tomato, shown as the number of eggs laid per female per day alive (y-axis) across three experimental treatments: bean mites (Control), tomato mites (Tomato), and tomato mites reverted to bean for two generations (Tomato reverted). Different letters above boxplots represent significant differences between treatments within each host plant. B) Reproductive performance of females from the same experimental treatments as in A, assessed in wildtype tomato cv. Castlemart, a tomato mutant impaired in jasmonic acid-related defences (Def1), and on this mutant tomato but rescued with exogenous application of jasmonic acid and isoleucine (Def1+JA). Boxplots show the median and the quartiles of the data, while dots show each individual female belonging to three replicates from the experimental evolution assay (legend in A).

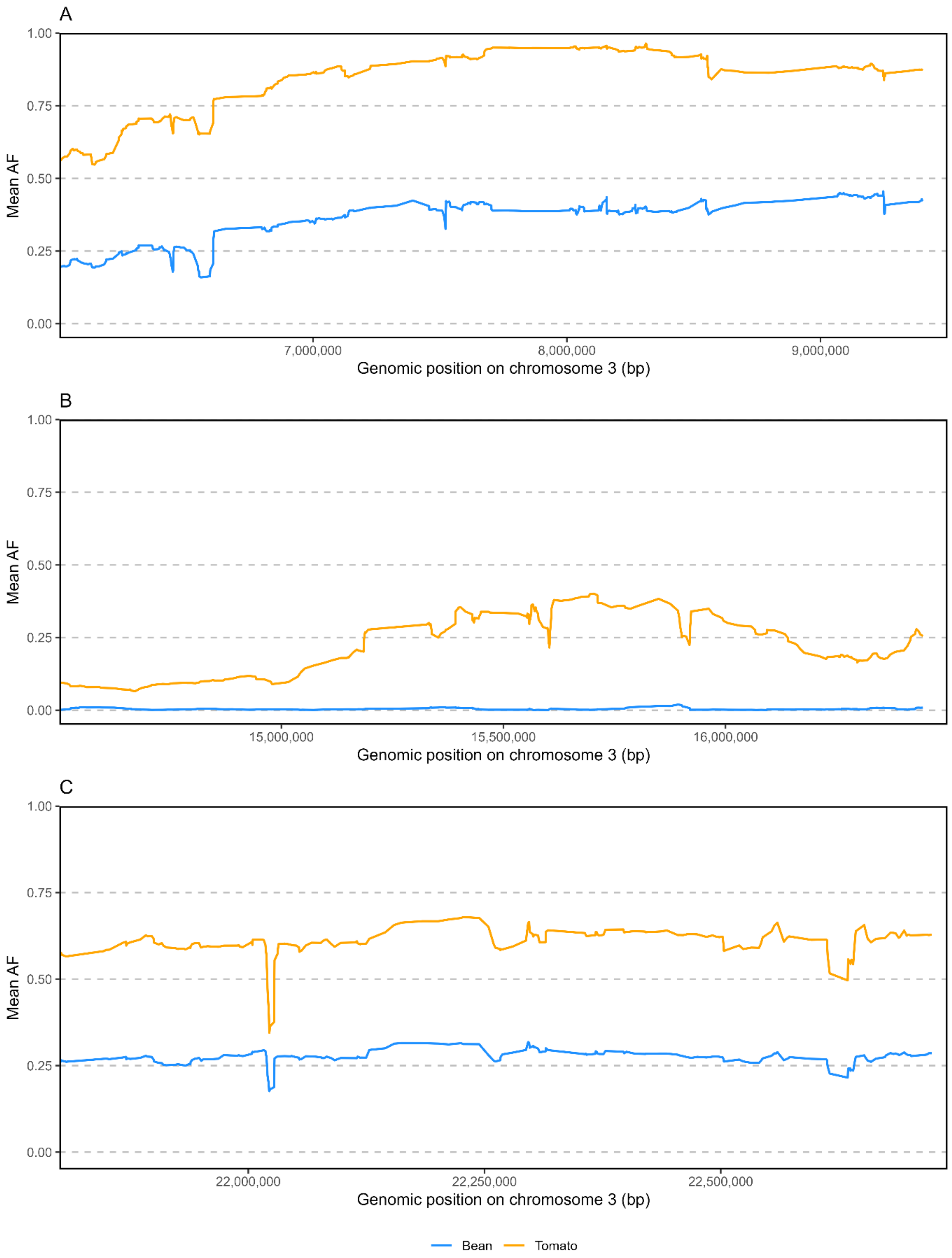


**Supplementary Figure 6. Mean allele frequency of unique parental markers SNPs in bean-control and tomato-selected replicates.** Mean allele frequency in sliding windows of 50 SNPs with a step size of 5 SNPs for (A) homozygous PL marker SNPs in the QTL1 region (B) homo- and heterozygous NL marker SNPs in the QTL2 region (C) homo– and heterozygous PL marker SNPs in the QTL3 region.

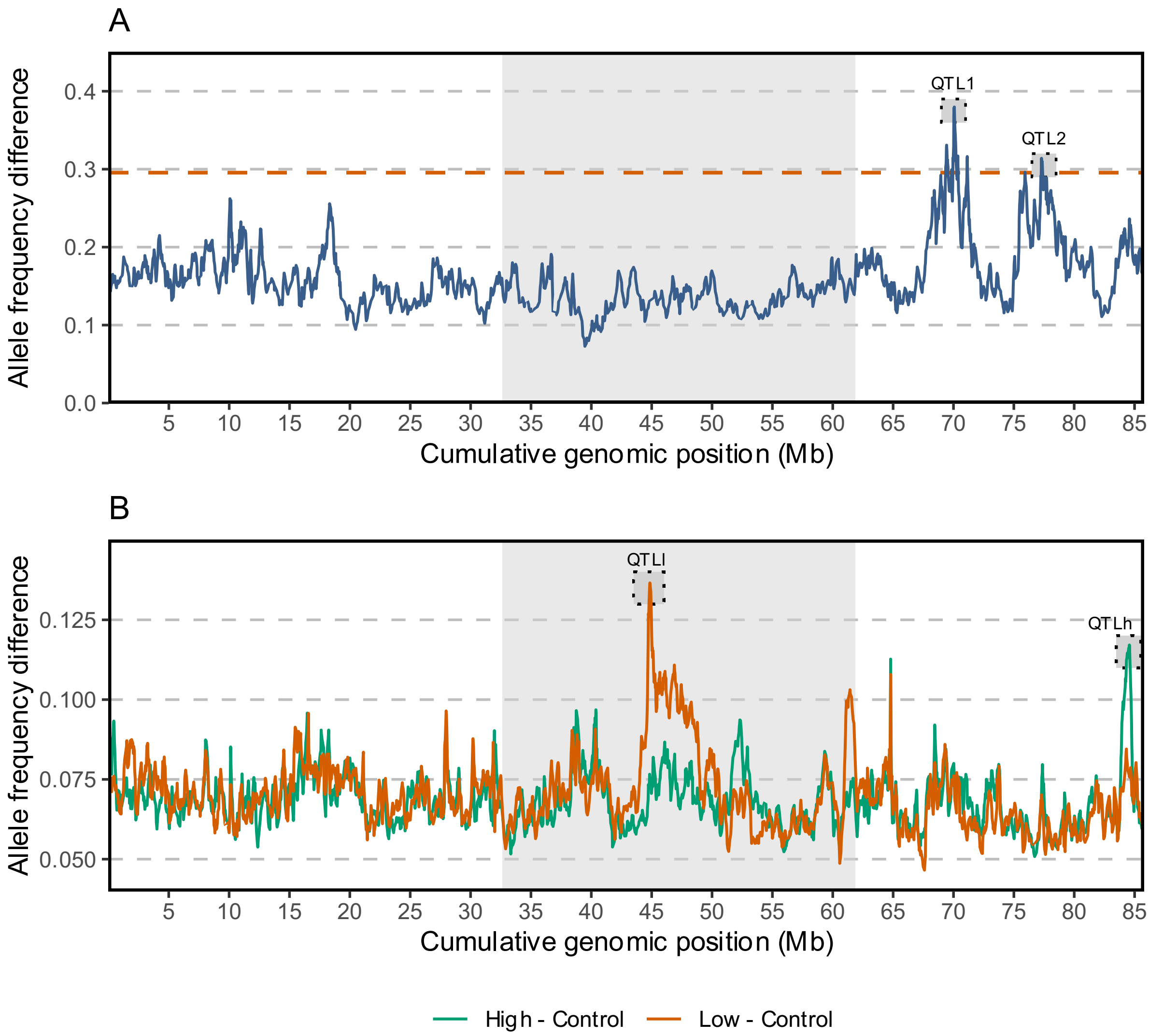


**Supplementary Figure 7. Mean allele frequency differences in sliding windows between bulks as assessed by standard BSA-script.** (A) Mean allele frequency difference between tomato-adapted and bean-adapted populations in 150kb sliding windows with a step size of 15 kb. The red dashed line denotes the significance threshold determined by a permutation approach (see Supplementary Note 3). (B) Mean allele frequency difference between the high performing and control bulk (green) and between the low performing and control bulk (red) in 250kb sliding windows with a step size of 25 kb. No significance threshold could be calculated due to the lack of replicates required for permutations in the BSA script.

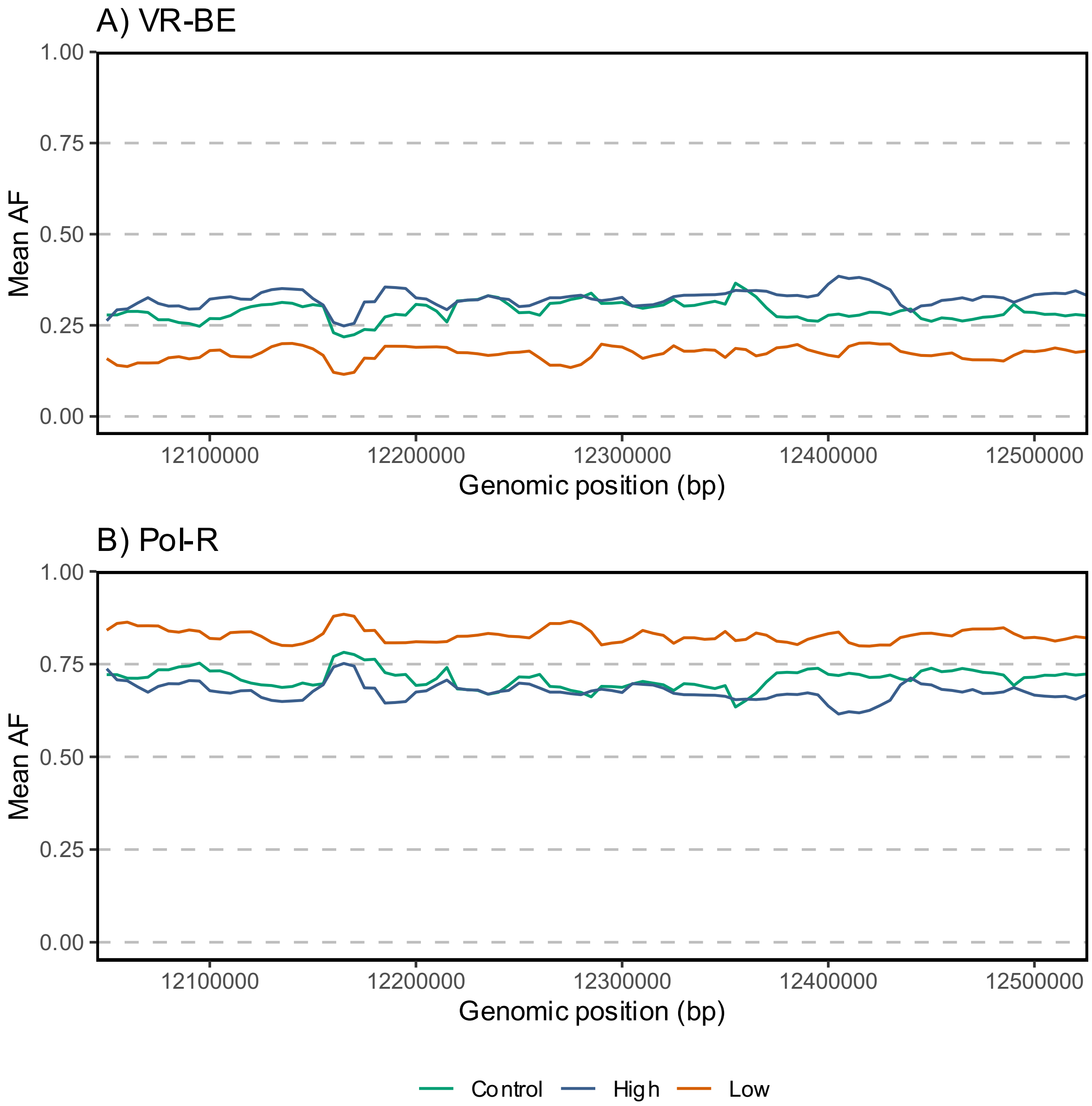


**Supplementary Figure 8. Mean allele frequency of unique parental markers SNPs in the pools.** Mean allele frequency in sliding windows of 25kb with a step size of 5kb for homozygous VR-BE (A) and Pol-R (B) marker SNPs in the QTLl region for the 3 different pools: unselected (control), high-performing and low-performing pool.
